# Environment-dependent selection impacts heritable developmental stability and trait canalization in rice

**DOI:** 10.1101/2025.06.13.659410

**Authors:** Taryn S. Dunivant, Irina Ćalić, Conor Gilligan, Zoé Joly-Lopez, Jae Young Choi, Mignon Natividad, Carlo Leo U. Cabral, Rolando O. Torres, Georgina Vergara, Steven J. Franks, Amelia Henry, Michael D. Purugganan, Simon C. Groen

## Abstract

Canalization, or the maintenance of trait values regardless of environmental or genetic variability, is fundamentally important for maintaining developmental stability. While this concept was described decades ago, we still know relatively little about how canalization is influenced by environmental stress, how it is shaped by natural selection, and the genetic underpinnings of canalization. In this study, we examined natural selection on microenvironmental canalization in rice (*Oryza sativa*) in wet and dry field conditions. We measured developmental stability in genetically identical replicates obtained from geographically widespread Indica and Japonica rice accessions, providing precise estimates of canalization in thousands of plants. We found that drought stress decreased canalization, showing that stress can increase instability. We also found evidence that canalization can evolve, given that canalization of several traits was heritable and under selection. We further uncovered specific genes underlying canalization, with genetic mapping and functional genetic experiments showing that the bZIP transcription factor-encoding gene *OsTGA5*/*rTGA2.3*, which is part of a module that balances stress response and plant growth, regulates canalization of several traits in an environment-dependent manner. At a genome-wide scale, canalization was associated with lower gene expression stochasticity at an earlier life stage, indicating that expression variation can reduce canalization and increase instability. Trait canalization was also positively correlated to temperature at accessions’ source environments, suggesting that selection on canalization can vary among environments. Overall, our study provides novel insights into the molecular genetic basis of environmental differences in developmental stability and how it might be shaped by selection.

**Significance:** Mechanisms stabilizing organismal development in response to genetic mutations or environmental stressors (canalization) have been reported for numerous animals and plants, but their underlying genetic basis and whether they may be shaped by selection remain unclear. Here, we report patterns of drought-induced trait decanalization in populations of rice (*Oryza sativa*) grown in field environments. We determined that trait canalization is heritable and can evolve separately from trait means. We identified the gene *OsTGA5*, part of a regulatory module shaping trade-offs between growth and stress responses, as impacting trait canalization. Plants with less noisy gene expression and evolving in warmer environments display greater developmental stability, contributing to the notion that canalization in rice may be adaptive.

## Introduction

The development of an organism is an extremely complex process involving the coordinated regulation of thousands of genes, transcription factors and other processes. Furthermore, development needs to be robust to genetic and environmental variation. But even genetically identical individuals developing in the same environment can show phenotypic differences because of developmental noise (Debat and David, 2001; Schlichting and Pigliucci, 1998). This developmental noise can be caused by developmental instability and microenvironmental variation, leading to unique trait variations such as differences in fingerprints between monozygotic twins (Glover et al., 2023). The specific mechanisms that drive developmental noise include variation in the timing and location of molecular interactions, which may be due to stochasticity in gene expression, random fluctuations in chemical or physical signaling, random partitioning of cytoplasmic components during cell division, and microenvironmental effects, both internal and external to the organism (Forde, 2009; Lachowiec et al., 2016; Viney and Reece, 2013). In some cases, developmental noise can have a genetic basis and be caused by such processes as spontaneous mutations, epigenetic changes in somatic tissues, epistasis, and genome heterozygosity (Forde, 2009; Lachowiec et al., 2016; Viney and Reece, 2013). Environmental stressors may impinge on the mechanisms that drive developmental noise, which could lead to environment-induced enhancement of phenotypic variability, i.e., stress-induced decanalization (Boukhibar and Barkoulas, 2016; Flatt, 2005; Hallgrimsson et al., 2019).

Environmental canalization, as proposed by Waddington and Schmalhausen in the 1940s, acts as a counterbalance to developmental noise by providing robustness. Canalization refers to the ability of organisms to produce consistent phenotypes despite variability in their environment or genotype. Canalization is the opposite of phenotypic plasticity, which is the ability of the same genotype to produce different phenotypes in different environments (Debat and David, 2001; Schlichting and Pigliucci, 1998). The buffering capacity of canalization helps maintain developmental stability and reduces phenotypic variation (Debat and David, 2001; Lachowiec et al., 2016). In this perspective, canalization is adaptive and expected to be under selection when it helps organisms maintain consistency in development across variable environments. But even if canalization is adaptive, it can have important consequences for evolvability and can result in either positive or negative effects on the ability of organisms to respond to environmental challenges. On one hand, canalization may allow for the accumulation of phenotypically cryptic genetic variation, which can be released after “decanalizing” events, allowing selection to act on newly expressed phenotypes (Flatt, 2005). On the other hand, canalization may temporarily constrain phenotypic evolution by keeping genetic variation phenotypically hidden (Flatt, 2005).

The balance between developmental noise and canalization may therefore influence the direction and rate of phenotypic evolution of populations (Flatt, 2005). Understanding the genetic basis of canalization and how canalization is shaped by selection could provide insights into the maintenance of genetic variation for phenotypes and the evolution of developmental systems, with implications for biomedicine and food production (Bruijning et al., 2020). For example, canalization potentially contributes to missing heritability in genomic predictions of medically and agriculturally relevant phenotypes (Hallgrimsson et al., 2019).

Many prior studies have examined canalization in model organisms such as *Caenorhabditis elegans* (Diaz and Viney, 2014); *Drosophila melanogaster* (Ayroles et al., 2015; Chen et al., 2015; Chen and Wagner, 2012; Dworkin, 2005; Lack et al., 2016; Morgante et al., 2015; Wang et al., 2017), and *Arabidopsis thaliana* (Hall et al., 2007; Jimenez-Gomez et al., 2011; Joseph et al., 2015) as well as in non-model organisms such as the leopard gecko *Eublepharis macularius* (Kiskowski et al., 2019). These studies were mostly conducted in controlled conditions, although one study of the great tit *Parus major* measured selection on canalization of fledgling weight in natural populations (Mulder et al., 2016). Some prior studies have investigated the genetic basis of canalization. Interestingly, the gene for the heat shock protein Hsp90 was identified as a candidate gene influencing canalization in the widely divergent species *D. melanogaster* (Chen and Wagner, 2012; Rutherford and Lindquist, 1998) and *A. thaliana* (Queitsch et al., 2002). This gene buffers against mutational variation, but it does not fully account for environmental canalization, suggesting the involvement of other genes (Milton et al., 2003). Because plants have a number of advantages for examining the genetic basis of and patterns of selection on trait canalization, they have been the focus of several studies, including ones on tomato (Alseekh et al., 2017; Fisher et al., 2017) and maize (Li et al., 2020). Some of these studies observed within-individual variation in fruit or seed size in species such as the hawthorn *Crataegus monogyna* (Sobral et al., 2014, 2019), spurge creeper *Dalechampia scandens* (Pélabon et al., 2021), and bread wheat *Triticum aestivum* (Beral et al., 2020). While these studies provide important information on canalization and its genetic basis, we still lack an integrative view of how the environment influences trait canalization, what the heritability and genetic basis of trait canalization is in different environments, how selection on canalization may differ between environments, and which environmental factors might shape the evolution of trait canalization.

To help address this need, we examined the genetic basis of and selection on microenvironmental canalization of 11 reproductive-stage traits in a diverse panel of rice accessions grown in wet and dry field conditions (Groen et al., 2020). Prior work on this same experiment characterized the strength and pattern of selection on gene expression (Groen et al., 2020), examined constraints to evolution caused by genetic architecture (Ćalić et al., 2022), and analyzed selection on macroenvironmental plasticity of gene expression (Hamann et al., 2024). The current study, which builds on this prior work, takes advantage of the fact that clonal replicates of hundreds of genotypes were planted in contrasting experimental conditions in the field, allowing a unique opportunity to examine canalization. Using this system, we obtained information on the genetic basis of canalization, including identifying a locus (*OsTGA5*/*rTGA2.3*) that may influence developmental stability in an environment-dependent manner. We also examined the strength and pattern of selection on trait canalization in a wet environment and under drought stress. Finally, we investigated if gene expression stochasticity and climatic factors at accessions’ source environments influence levels of trait canalization.

## Results

### Drought Stress Induces Trait Decanalization

From our field experiment, we quantified microenvironmental canalization within 2-m-wide row plots for five plants per plot, with each plant spaced 0.2 m apart from any neighboring plants (Fig. S1a), considering 11 reproductive-stage morphological traits [plant height (HGT), shoot dry weight (vegetative parts; SDW), tiller number (TNR), days to panicle maturity (DPM), panicle number (PNR), panicle length (PNL), rachis dry weight (RDW), primary panicle branch number (PBN), secondary panicle branch number (SBN), spikelet number (SPN), and hundred-grain weight (HGW)] in wet and dry conditions (Data S1-4). We considered 93 Indica varietal group accessions (including indica and *circum*-aus) and 50 Japonica varietal group accessions (including temperate and tropical japonica and *circum*-basmati) as separate populations to account for population structure (Fig. S1b; Groen et al., 2020). For each trait, we calculated within-genotype variation using the log-transformed Levene’s Statistic (LS) as our estimate for canalization (Hall et al., 2007). We observed a significant effect of drought on trait means (two-way ANOVA, p≤0.0444) as well as significant drought-induced decanalization for all traits (two-way ANOVA, p≤0.00414) in Indica (Fig. 1a, Figs. S2-3) and Japonica accessions (Figs. S4-5) except for TNR among Japonica accessions, for which the environmental effect on canalization was marginally non-significant (two-way ANOVA, p=0.06345). These results show that, as expected, stressful dry field conditions affect trait means and engender a decrease in canalization of nearly all traits.

**Fig. 1.**
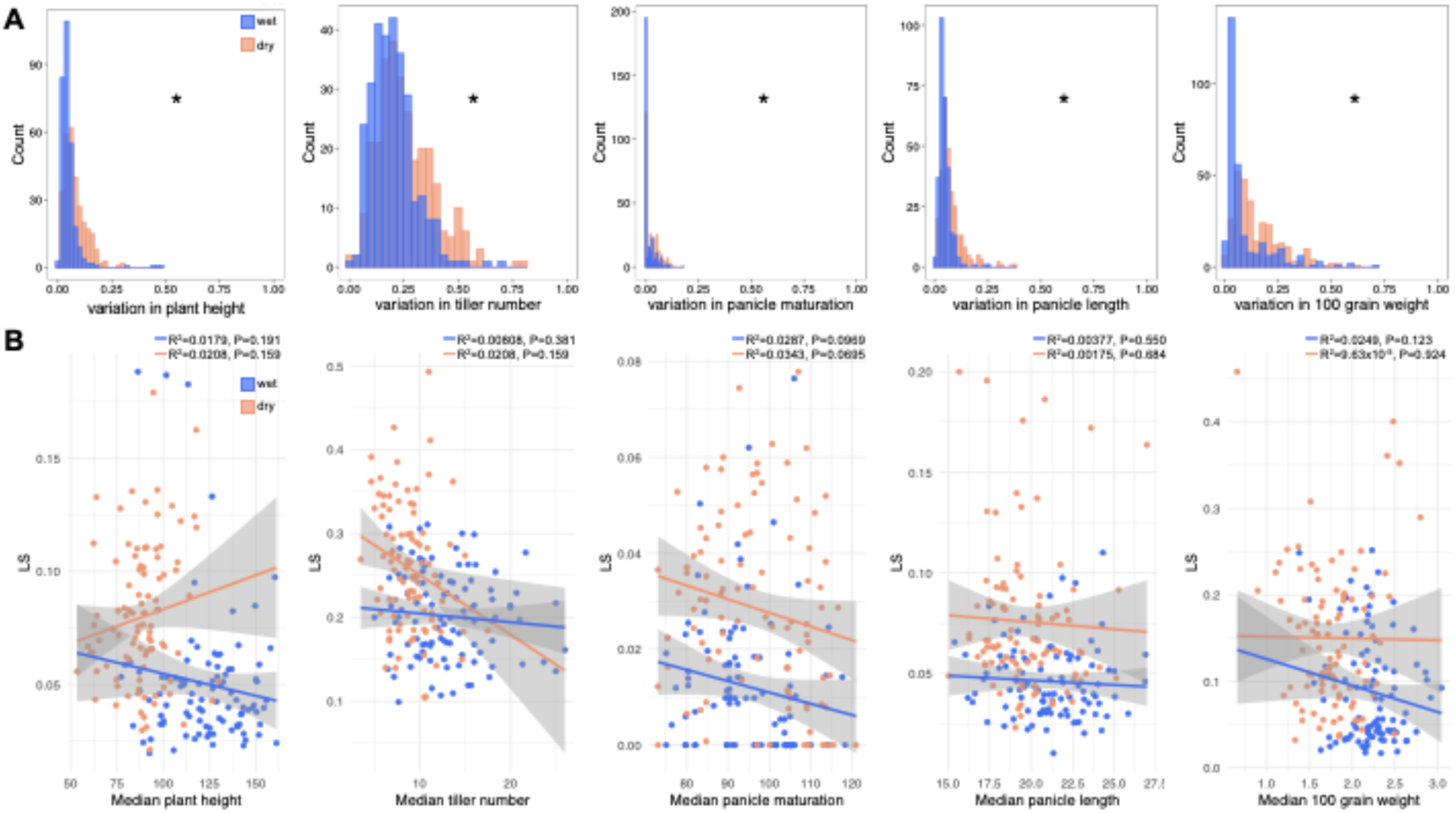
Drought induces trait decanalization in Indica rice accessions. (*A*) Frequency distributions for trait canalization (as measured through micro-environmental variation using Levene’s Statistic) for five morphological traits in Indica accessions (n=93). From a two-way ANOVA, environment was significant for all traits (p<0.05). (*B*) Genotypic correlations between trait means and trait canalization (as measured through micro-environmental variation) for five morphological traits in Indica accessions. No significant Pearson product-moment correlations were identified (p>0.05). Blue indicates wet conditions and orange indicates dry conditions.

### Trait Canalization Shows Varying Heritability and May Evolve Separately from Trait Means

Next, we estimated broad-sense heritabilities of microenvironmental variations in traits (as measured by LS) and trait means for the Indica and Japonica populations in wet and dry conditions (Table 1, Table S1-2). We observed significant heritability for all trait means in both populations as well as for canalization of DPM in wet conditions for Indica, of HGW and PNL for Indica under drought, of PNL for Japonica in the wet and dry field environments, and of DPM and HGW for Japonica under drought (two-way ANOVA, p≤0.0433). This suggests that canalization of these traits may evolve in the Indica and Japonica populations under certain environmental conditions. We also found that trait means and trait canalization are expected to evolve independently, given that there were no significant genetic correlations between trait means and microenvironmental variations in traits for the Indica and Japonica populations in either wet or dry conditions (Pearson’s r, p>0.05; Fig. 1b, Figs. S6-7).

**Table 1.** Broad-sense heritabilities (*H^2^*) for trait means and canalization as well as selection gradients (*β*) for trait canalization in the Indica population.

| | $H^2$ GxE | | $\beta$ wet | | $\beta$ dry | |
| --- | --- | --- | --- | --- | --- | --- |
|  | Median | LS | Estimate | Std Error | Estimate | Std Error |
| **HGT** | <u>0.7286</u> | 0.07245 | -0.0415 | 0.0377 | 0.0939 | 0.0771 |
| **TNR** | <u>0.6773</u> | 0.0000 | -0.053 | 0.0322 | 0.095 | 0.0799 |
| **SDW** | <u>0.6314</u> | 0.0000 | **<u>-0.0987</u>** | 0.0343 | 0.0017 | 0.0685 |
| **DPM** | <u>0.8511</u> | **<u>0.2190</u>**<sup>+</sup> | -0.0263 | 0.0415 | -0.0668 | 0.0442 |
| **PNR** | <u>0.5575</u> | 0.0000 | -0.0038 | 0.0325 | -0.0635 | 0.0639 |
| **PNL** | <u>0.7469</u> | 0.1634* | **<u>-0.1356</u>** | 0.0375 | **<u>-0.2218</u>** | 0.0736 |
| **RDW** | <u>0.3299</u> | 3.664x10<sup>-9</sup> | **<u>-0.0611</u>** | 0.0258 | -0.1325 | 0.0744 |
| **PBN** | <u>0.6408</u> | 0.0000 | -0.0226 | 0.0403 | **<u>-0.1496</u>** | 0.07 |
| **SBN** | <u>0.5347</u> | 0.1437 | 0.0426 | 0.0396 | -0.1333 | 0.073 |
| **SPN** | <u>0.3810</u> | 0.0000 | -0.0308 | 0.021 | -0.1247 | 0.072 |
| **HGW** | <u>0.7395</u> | **<u>0.04422</u>**<sup>*</sup> | -0.0059 | 0.0383 | **<u>-0.4445</u>** | 0.0604 |

Table 1. Broad-sense heritabilities ( $H^2$ ) for trait medians and canalization, and selection gradients ( $\beta$ ) for trait canalization in the Indica population.
| | $H^2$ GxE | | $\beta$ wet | | $\beta$ dry | |
| --- | --- | --- | --- | --- | --- | --- |
|  | Median | LS | Estimate | Std Error | Estimate | Std Error |
| **HGT** | <u>0.7286</u> | 0.07245 | -0.0415 | 0.0377 | 0.0939 | 0.0771 |
| **TNR** | <u>0.6773</u> | 0.0000 | -0.053 | 0.0322 | 0.095 | 0.0799 |
| **SDW** | <u>0.6314</u> | 0.0000 | **<u>-0.0987</u>** | 0.0343 | 0.0017 | 0.0685 |
| **DPM** | <u>0.8511</u> | **<u>0.2190</u>**<sup>+</sup> | -0.0263 | 0.0415 | -0.0668 | 0.0442 |
| **PNR** | <u>0.5575</u> | 0.0000 | -0.0038 | 0.0325 | -0.0635 | 0.0639 |
| **PNL** | <u>0.7469</u> | 0.1634* | **<u>-0.1356</u>** | 0.0375 | **<u>-0.2218</u>** | 0.0736 |
| **RDW** | <u>0.3299</u> | 3.664x10<sup>-9</sup> | **<u>-0.0611</u>** | 0.0258 | -0.1325 | 0.0744 |
| **PBN** | <u>0.6408</u> | 0.0000 | -0.0226 | 0.0403 | **<u>-0.1496</u>** | 0.07 |
| **SBN** | <u>0.5347</u> | 0.1437 | 0.0426 | 0.0396 | -0.1333 | 0.073 |
| **SPN** | <u>0.3810</u> | 0.0000 | -0.0308 | 0.021 | -0.1247 | 0.072 |
| **HGW** | <u>0.7395</u> | **<u>0.04422</u>** * | -0.0059 | 0.0383 | **<u>-0.4445</u>** | 0.0604 |

$H^2$ significance by environment: <sup>+</sup> wet, \* dry.

### Selection Influences Trait Canalization and Varies by Environment

To assess if selection might act on trait canalization, we performed multivariate genotypic selection analyses (Lande and Arnold 1983, Rausher 1992) on microenvironmental variations in traits while accounting for trait means. For both Indica and Japonica, selection tended to favor larger phenotypic values in wet conditions, with more variable patterns of selection under drought (Table 1; Table S3). For Indica, we observed significant positive selection on trait canalization (as indicated by a negative selection gradient *β* on microenvironmental trait variation) for PNL, RDW, and SDW in wet conditions as well as for HGW, PBN, and PNL under drought (Table 1). For Japonica, we found significant positive selection on trait canalization for SPN in the wet field environment as well as for HGW and SBN under drought (Table S4). By contrast, the only trait for which we observed negative selection on trait canalization (as indicated by a positive selection gradient *β* on microenvironmental trait variation) was for SPN among Japonica accessions under drought (Table S4). Since canalization of HGW and PNL in Indica and HGW in Japonica showed significant heritability under drought, and since heritability of PNL is also substantial for Indica rice in wet conditions (Table 1, Table S2-3), positive selection on the canalization of these traits has the potential to change the microevolutionary dynamics of canalization.

### OsTGA5 May Regulate Trait Canalization

In the Indica population under drought, we identified two loci associated with canalization of panicle length (PNL) through genome-wide association mapping (Fig. 2a; Data S5). The first was tagged by a SNP on chromosome 10 that is significantly associated with PNL and is located within a gene encoding an F-box domain-containing protein, *OsFbox536* (Os10g0145100; Fig. 2b). This gene is part of an abiotic stress-responsive gene coexpression module containing genes involved in RNA splicing (Krishnan et al., 2017). Furthermore, this locus contains a cluster of genes encoding F-box domain-containing proteins and has previously been associated with variation in panicle length of rice plants grown under abiotic stress (Lekklar et al., 2019).

**Fig. 2.**
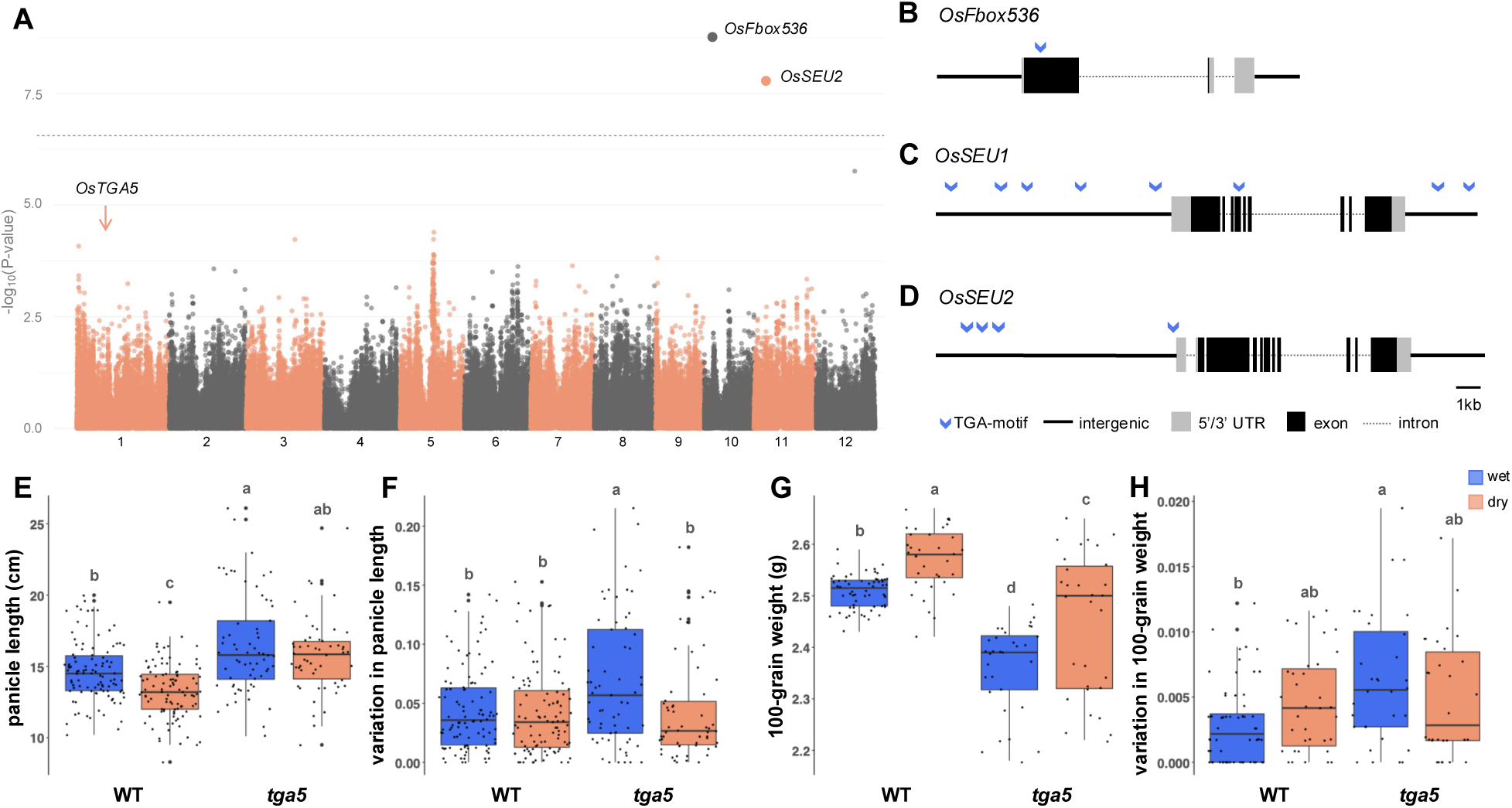
OsTGA5 regulates trait canalization. (A) Manhattan plot showing the results of genome-wide association mapping performed using a multi-locus linear mixed model (MLMM) with 179,634 single-nucleotide polymorphisms (SNPs) as markers. SNPs in/near *OsFbox536* and *OsSEU2* were significantly associated with panicle length (PNL) canalization for the Indica population under drought after p-value correction using a Bonferroni threshold of p<2.78 × 10^−7^. The location of *OsTGA5* is indicated with an arrow. (*B-D*) TGA transcription factor-binding elements (TGA motif: TGACGT) in and around *OsFbox536* (*B*) as well as *OsSEU1* and -*2* (*C* and *D*, respectively). (*E-H*) Final measurements of trait means and canalization (as measured through micro-environmental variation) for PNL (*E-F*) and hundred-grain weight (HGW, *G-H*) of wild-type Nipponbare and *tga5* loss-of-function mutant plants in wet and dry conditions showed significant genotype, environment, and genotype-by-environment interaction effects (two-way ANOVA, p<0.05), except for mean PNL, where only the former two effects were significant. Blue indicates wet conditions and orange indicates dry conditions.

Another significant SNP for PNL, on chromosome 11, is located 8,010 bp downstream of the gene *OsSEU2* (Os11g0207100; Fig. 2c). This gene and its family member *OsSEU1*, located in tandem (Os11g0207000), are co-orthologs of Arabidopsis *AtSEU* and are involved in various developmental processes (Tanaka et al., 2017). Expression of both *OsSEU1* and *-2* is strongly co-regulated with expression of *OsTGA5*/*rTGA2.3*, which encodes a bZIP transcription factor (TF) that has been identified as a regulator of stress responses (Niu et al., 2022; Sato et al., 2013). Furthermore, we identified binding sites for bZIP TFs in the promoters and exonic regions of *OsFbox536* as well as of *OsSEU1* and *-2* (Fig. 2b-d, Table S5). Given the potential role of OsTGA5 in regulating all three of these genes, we decided to test if OsTGA5 is involved in regulating trait canalization in rice (Data S6). Knocking out *OsTGA5* function using CRISPR gene editing in the genetically manipulable Japonica accession Nipponbare led to an increase in panicle length in wet and dry greenhouse conditions (two-way ANOVA; *F_genotype_*=57.454, p=3.87 × 10^−13^; *F_environment_*=17.878, p=3.09 × 10^−5^; Fig. 2e, Fig. S8) and altered canalization of panicle length in a genotype-, environment-, and genotype-by-environment-dependent manner (two-way ANOVA; *F_genotype_*=8.659, p=0.00349; *F_environment_*=5.869, p=0.01597; *F_G×E_*=5.611, p=0.01845; Fig. 2f). Other traits were similarly impacted by knocking out *OsTGA5* (Fig. 2g,h, Fig. S8-11), showing that this gene has the potential to regulate trait canalization.

### Genome-wide Gene Expression Stochasticity Influences Trait Canalization

To determine if noisiness of gene expression earlier in life might impact canalization of traits later in life, we obtained measures of genome-wide gene expression stochasticity (Groen et al., 2020; Jimenez-Gomez et al., 2011). Continuing to focus on the larger Indica population, we identified three traits whose levels of canalization were negatively correlated with global stochasticity in gene expression (as defined by positive Pearson correlations between expression stochasticity and trait variation) in wet conditions: PNR, RDW, and SPN (Pearson’s *r*≥0.206, p≤0.0478; Fig. 3a; Data S7). Drought stress caused significant decanalization of global gene expression stochasticity (Student’s *t*=2.5768, p=0.0108; Fig. 3b), similar to what we observed for the morphological traits (Fig. 1). However, despite a significant correlation of global gene expression stochasticity in the wet and dry field environments (Pearson’s *r*=0.602, p=1.70×10^−10^), the significance of correlations between expression stochasticity and trait canalization disappeared under drought stress (Pearson’s *r*, p>0.05).

**Fig. 3.**
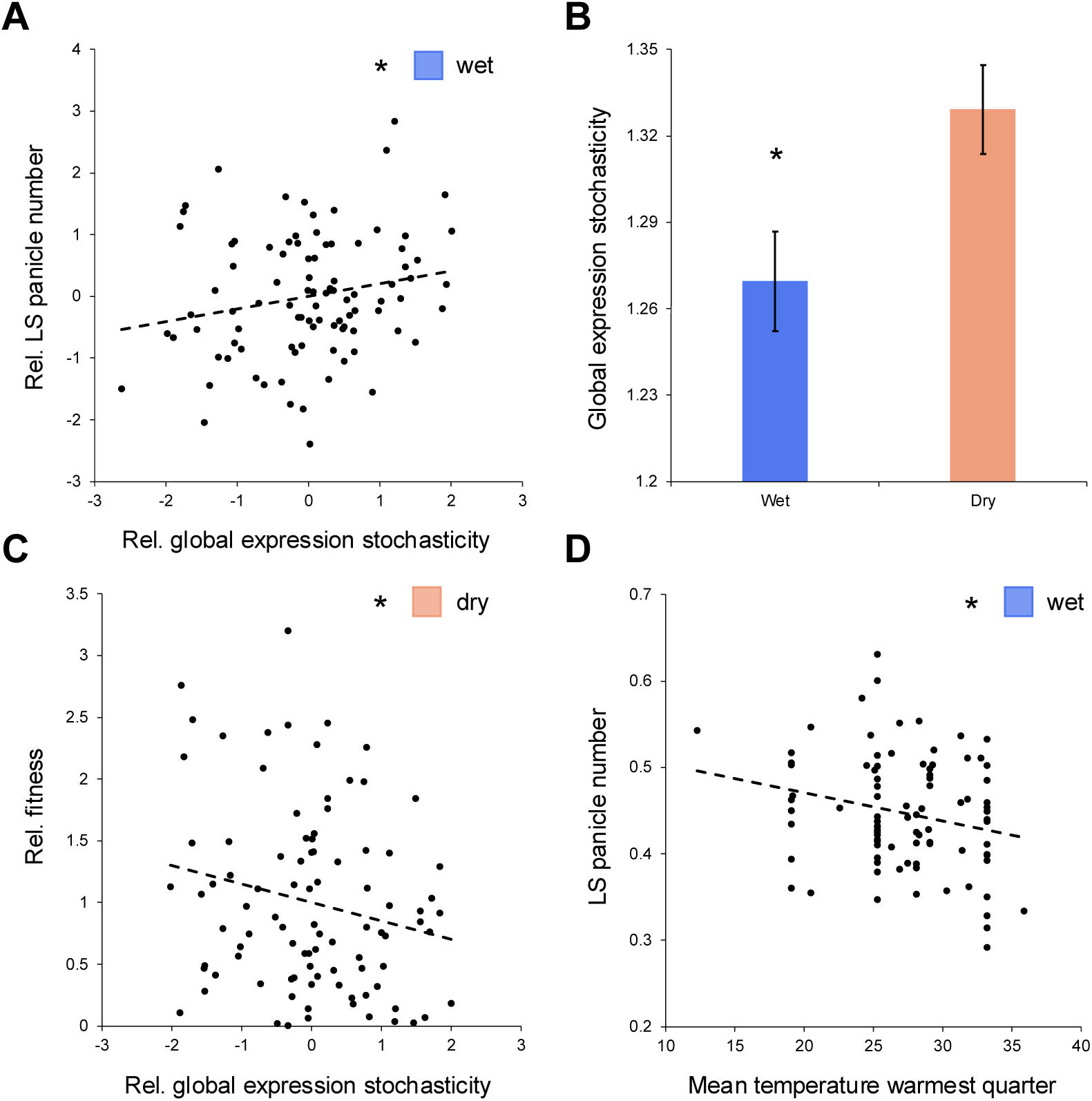
Global gene expression stochasticity and environment-of-origin influence trait canalization in Indica rice. (*A*) Genome-wide gene expression stochasticity showed a significantly positive correlation with micro-environmental plasticity (as determined by the log-transformed Levene’s Statistic) of panicle number (PNR) in the Indica population (n=93) in wet conditions: i.e., a negative correlation with canalization of this trait (Pearson’s *r*=0.206, p=0.0478). (*B*) Global gene expression stochasticity is higher under drought than in wet conditions (Student’s *t*=2.5768, p=0.0108). (*C*) There was weak selection against global gene expression stochasticity in the Indica population under drought (linear selection gradient *β*=0.15, one-tailed p=0.026). (*D*) The temperature of the warmest quarter showed a significantly negative Pearson product-moment correlation with micro-environmental plasticity of PNR in wet conditions: i.e., a positive correlation with canalization of this trait (Pearson’s *r*=-0.231, p=0.0258).

These findings imply that accessions with higher genome-wide gene expression stochasticity during development may later show lower levels of canalization for at least some traits in relatively stress-free wet field conditions. Congruent with the selection for canalization of RDW and other traits as well as the mostly negative or non-significant correlations between trait canalization and global gene expression stochasticity in at least the wet environment (*χ^2^_wet_*=4.455, p=0.0348), we observed weak negative directional selection on expression stochasticity in wet and dry conditions (*β_wet_*=-0.068, one-tailed p=0.0455; *β_dry_*=-0.15, one-tailed p=0.026; Fig. 3c). In addition, the mostly negative or non-significant correlations between trait means and global gene expression stochasticity further suggests that noise in gene expression might be costly (*χ^2^_wet_* and *χ^2^_dry_*=7.364, p=0.0067).

### Source Environment Influences Trait Canalization

Finally, we aimed to explore if the prevailing climate in the source environments of Indica rice accessions might shape patterns of trait canalization. We found that canalization of PNR, SDW, and TNR correlated positively with the bioclimatic variable of mean temperature in the warmest quarter of the year at the accessions’ source environments (as indicated by negative Pearson correlations between temperature in environments-of-origin and trait variation) for plants growing in the wet and relatively hot conditions of the Philippines dry season (Pearson’s *r*≤-0.224, p≤0.031; Fig. 3d; Data S7), but not when drought stress was added to these conditions. These observations are consistent with local adaptation, with accessions from warmer and more stressful environments having evolved a stronger capacity for trait canalization than accessions from cooler climates.

## Discussion

In this study of multiple genotypes of field-grown rice plants, we show that drought stress generally reduces trait canalization, that canalization shows variable heritability across traits, and that trait canalization may evolve separately from trait means. We find that natural selection can impact canalization of traits. Since canalization of many traits showed substantial heritability, selection further has the potential to change the microevolutionary dynamics of canalization.

Combining our measures of canalization for reproductive-stage traits with estimates of genome-wide stochasticity in gene expression, we identified neutral to negative correlations between gene expression noise and trait canalization (Groen et al., 2020). Similar links between gene expression stochasticity and developmental stability have previously been observed in the nine-banded armadillo *Dasypus novemcinctus* (Ballouz et al., 2023) and in *A. thaliana* (Cortijo et al., 2019; Jimenez-Gomez et al., 2011). The environment-dependent decanalization of genome-wide gene expression we observed is in keeping with what has previously been found in *D. melanogaster* (Chen et al., 2015). Stochasticity in gene expression may contribute to environment-induced regulatory decoherence of gene expression, a phenomenon that has been described in the context of immune challenges in humans and salt stress in rice (Gupta et al., 2025; Lea et al., 2019). Future studies could bring more insight into the molecular mechanisms that determine canalization of gene expression. Candidate mechanisms include cell-to-cell variability in DNA methylation as well as the occurrence of somatic mutations and concomitant effects on gene expression variability (Cruzan et al., 2022; Goeldel and Johannes, 2023; Wang et al., 2019).

We were able to uncover part of the genetic underpinnings of canalization of certain morphological traits, in particular panicle length (PNL). Our mapping efforts revealed an association of PNL canalization with genetic variation in *OsSEU1* and *-2*, which are coexpressed tightly with the bZIP TF-encoding gene *OsTGA5*/*rTGA2.3* (Sato et al., 2013). The OsSEU1 and −2 proteins interact with OsLUGL, forming transcriptional co-regulatory complexes that control identity and development of organs, including the panicle, by interacting with MADS TFs. These MADS TFs include OsMADS18 (Yang et al., 2019), which we and others previously identified as a regulator of rice responses to drought, including changes in panicle size (Groen et al., 2020; Yin et al., 2019). OsTGA5/rTGA2.3 is further a known interacting partner of OsNPR1, a major regulator of the balance between plant growth and responses to environmental stressors (Xu et al., 2017). Overexpression of *OsNPR1* is known to lead to restrained growth and development phenotypes, which can be restored by silencing expression of the Gretchen Hagen 3 (GH3) acyl acid amido synthetase OsGH3-8 in parallel. Reducing OsGH3-8 activity can partially restore levels of auxin and largely rescue the developmental phenotype of *OsNPR1*-overexpressing plants without affecting responses to changes in abiotic environmental factors (Li et al., 2016). Interestingly, OsGH3-8 is also an important target of OsLUGL-OsSEU and MADS TF regulatory complexes (Yang et al., 2019). Since altered expression of *OsGH3-8* leads to changes in panicle lengthening via changes in auxin signaling (Yadav et al., 2011), OsGH3-8 might be a central player that OsTGA5/rTGA2.3, and the regulators it is coexpressed with, impinge on.

It may be a fruitful endeavor to investigate further post-translational regulatory mechanisms that could influence developmental stability in the face of environmental stressors. OsSEU and other SEUSS proteins contain intrinsically disordered regions and likely alter plant growth and development during abiotic stress by condensating and recruiting transcriptional activators (Wang et al., 2022). These activators could potentially include OsTGA5/rTGA2.3 with which both OsSEU1 and −2 are strongly coexpresssed. OsTGA5/rTGA2.3 in turn shows evidence of post-translational regulation and may not only form homodimers but also heterodimerize with OsTGA2 and OsTGA3 (Niu et al., 2022).

One of the features distinguishing our field study is that we were able to observe that selection promoted canalization of both genome-wide gene expression and several morphological traits, with at least some of these traits showing a significant negative correlation between gene expression noise and trait canalization in the wet environment. Such links between organismal fitness and canalization of traits across levels of biological organization have been exceedingly difficult to measure. Indeed, we still have limited understanding of whether intra-genotypic phenotypic variability may be adaptive (Bruijning et al., 2020). One source of evidence that trait canalization may be adaptive in an agricultural environment is a set of studies showing that plants may start to forage for resources such as light and water inefficiently when growing in proximity to genetically identical or different neighboring plants in a suboptimal phenotypically plastic response to cues from the surrounding microenvironment. The ensuing microenvironment-induced trait decanalization is known as a “tragedy of the commons” and can to some extent be prevented through breeding crops according to phenotypically canalized ideotypes that do not engage in competition with neighboring plants (Anten and Vermeulen, 2016). A second source of evidence that trait canalization may be adaptive more generally is if traits show more variability for accessions at the edge of a species’ range, in environments that were colonized relatively recently. Here, we identified that canalization of several traits in wet conditions correlated positively with prevailing growing season temperatures at accessions’ source environments. In light of Asian rice’s domestication history, where it was not introduced into cooler, temperate environments until after having been established as a crop in tropical climes (Gutaker et al., 2020), this could suggest that accessions that currently hail from more temperate areas are more sensitive to trait decanalization when confronted with a stressful environment that is warmer and drier than what they have recently experienced. Analogous results have been obtained for *D. melanogaster*, where flies that had relatively recently migrated into a lower-temperature, high-altitude environment in Ethiopia were more sensitive to trait decanalization than flies from the ancestral range around Zambia (Lack et al., 2016). These results were interpreted as showing that decanalized development might represent a cost of adapting to a novel environment. In the future, it will be interesting to find out if such costs might play a role in adaptation of crop plants and other organisms to novel environments as well.

One trait for which we observed selection promoting canalization under drought in both the Indica and Japonica populations was grain size (HGW). Previous studies on the perennial vine *Dalechampia scandens* obtained evidence supporting the hypothesis that reduced canalization of seed size can be adaptive when environmental predictability is low (Pélabon et al., 2021). Studies with the perennial shrub hawthorn (*C. monogyna*), on the other hand, found that seed-dispersing birds and rodent seed predators exerted selection for less variable fruit and seed size in some environments (Sobral et al., 2014, 2019). Despite the reduced environmental predictability for rice plants in our dry field versus our wet field (plants experienced alternating cycles of drought followed by rewatering in the dry field but only stable well-watered conditions in the wet field), we observed selection for canalization of grain size in dry conditions. Interestingly, natural selection appears to shape canalization of this trait in a direction that is also desired by rice breeders, namely selection for increased uniformity (Anten and Vermeulen, 2016; Bruijning et al., 2020). Similar patterns were observed in field experiments with bread wheat (*T. aestivum*) grown across four different wet and dry environments, where grain size variance showed consistent negative correlations with an important fitness component, number of spikes per m^2^, which in turn was positively correlated with grain yield (Beral et al., 2020).

Overall, in this study, we found genetic evidence for links between trait canalization and fitness of Asian rice in two field environments, where microenvironmental differences are magnified over those arising in controlled environments. Trait canalization is a key developmental characteristic of living systems (Debat and David, 2001; Flatt, 2005), and elucidating its genetic basis as well as how it is influenced by selection is important for understanding how phenotypic stability evolves and may shape patterns of organismal development. We identified genes that may influence canalization levels across different environments and showed how selection may impact canalization for various traits. These findings lay the groundwork for further developing our understanding of the molecular genetic mechanisms and evolutionary processes that influence developmental stability.

## Materials and Methods

### Establishment of the Field Experiment

A detailed description of the field experiment, which was conducted during the 2016 dry season at the International Rice Research Institute in Los Baños (Laguna, Philippines), has been published previously (Groen et al., 2020). Briefly, 17-day-old seedlings of 220 accessions (Data S1), including two additionally replicated checks (IR64 and Sahod Ulan 1), were transplanted into two different experimental fields: one that remained flooded as a well-watered, wet paddy environment (UJ) and one within a rain-out shelter (UR) that was maintained flooded until 16 days after transplanting (DAT), when the field was drained upon stopping irrigation to initiate an intermittent drought stress treatment that was maintained until the end of the season by periodic rewatering through flooding at 36, 47, and 74 DAT. The seedlings were transplanted following an identical alpha lattice design in each field with triplicate plots per accession and each plot consisting of a single 2-m row of 10 isogenic seedlings. Rows were spaced 0.2 m apart and the within-row distance between seedlings was 0.2 m as well with one seedling transplanted per hill (Fig. S1a). To avoid induction of microenvironmental plasticity through biotic interactions, weeding was performed manually on a regular basis, while the insecticides Cymbush and Cartap were applied at 20 DAT, and Provado was applied at 23 and 43 DAT.

### Plant Harvesting and Processing

Individual panicles were harvested separately from each plant within the 4th, 6th, 7th, 8th, and 9th hills per plot as plants finished maturing (Fig. S1a). Remaining plant material was then harvested on an individual plant basis and processed by separating the panicles and tillers, for a total of 6,224 plants harvested individually (five plants per plot). Tillers were counted separately from each plant. The lengths of all panicles harvested were measured, and three panicles per plant were used to count the primary, secondary and tertiary branching of panicles. A seed counter (Hoffman Manufacturing Inc) was used to count filled, partially filled, and unfilled grains after sorting, with the exception of counting awned grains, which were processed manually. After oven-drying the grains at 45°C for 3 days, the weights of filled, partially filled, and unfilled grains were obtained for calculating HGW. The filled-grain-number data for the plants in the 4^th^ hill of each plot alone were previously published (Groen et al., 2020).

### Measuring Variation in Rice Populations

To obtain robust estimates of microenvironmental canalization and to allow comparisons of canalization across the wet and dry field environments, only accessions with non-zero fecundity for each of the three replicate plots per accession in each environment were included for downstream analyses. This filtering step left 93 accessions of the Indica varietal group (including indica and *circum*-aus accessions) that constituted one set of populations and 50 accessions of the Japonica varietal group (including japonica and *circum*-basmati accessions) that constituted a second set of populations (Data S2).

We performed further analyses for the Indica and Japonica populations separately to control for the major source of population structure in *O. sativa*. We estimated within-genotype variability by calculating the Levene’s Statistic (LS) for each trait per accession as a measure of microenvironmental canalization: *LS* = | log (*x*)*ij* – median (log (*Xj*)) |, where i denotes each replicate plant from accession j (Dworkin, 2005; Hall et al., 2007). We used the log-transformed LS based on the median as a standardized measure of within-genotype trait variability since this minimizes covariation between the LS and trait size (Dworkin, 2005; Hall et al., 2007). We calculated the LS for each replicate plot individually, which resulted in three replicate estimates of the LS per genotype-environment combination. We considered trait measures combined across the two environments and separately within each environment and processed them by ANOVA to partition phenotypic variation. For each trait, we fit mixed-effect general linear models for their plot median and LS values. The models included terms for accession or genotype (G) as a random factor, field environment (E) as a fixed factor, G × E interaction as a random factor for across-environment analysis only, block effects (B) accounting for the alpha lattice field layout, and error variance (ε). We tested if factors explained significant variance for each trait and its canalization using F-tests in the *lme4* package version 1.1 in *R* version 4.0.3.

We estimated broad-sense heritabilities for trait means (based on the median trait value per plot) and trait canalization (based on the LS per plot) as described previously (Groen et al., 2020; Hamann et al., 2024), with across-environment heritabilities estimated as *H^2^* = *σ^2^_G_* / ( *σ^2^_G_* + ( *σ^2^_GE_* / *e* ) + ( *σ^2^_E_* / *re* )) — *σ^2^_G_*, *σ^2^_GE_*, and *σ^2^_E_* are the among-genotype, G × E, and within-genotype variance components, respectively, *e* is the number of environments, and *r* is the number of replicates per environment — and within-environment heritabilities estimated as *H^2^* = *σ^2^_G_* / ( *σ^2^_G_* + *σ^2^_E_* ).

We determined genetic correlations between trait means and trait canalization as Pearson’s product-moment correlation coefficients using the cor function in R version 4.0.3.

### Estimating Genotypic Selection

We estimated selection on trait canalization through genotypic selection analysis (Rausher, 1992) by regressing averages for trait LS values per genotype against mean fitness per genotype within each environment (Hamann et al., 2024). We used relative fecundity fitness, measured as the average filled-grain number across replicate plots of each genotype within a field divided by the mean of all genotypes for that field, as our fitness measure. Genotypic trait means and trait canalization values were based on averaging the median trait and LS values per plot, respectively, for each genotype and were then standardized for all genotypes (mean=0, SD=1) within each environment.

To examine selection on trait canalization (selection on the degree to which a genotype maintains stable trait expression within an environment), we estimated *β* as the partial regression coefficient of relative fitness on the standardized genotypic mean trait value and standardized genotypic average trait canalization value (Hamann et al., 2024).

A positive partial regression coefficient of relative fitness on trait canalization value indicates selection for microenvironmental plasticity of a trait, while a negative coefficient indicates selection for trait canalization. Because the model includes both trait mean and canalization values, the partial regression coefficient for trait canalization reflects selection on canalization itself, separate from selection on the trait mean produced through canalization (Hamann et al., 2024).

### Genome-Wide Association Mapping of Canalization Genes

We conducted GWA mapping using the R package GAPIT version 3 by running a multi-locus linear mixed model (MLMM) for each trait mean and canalization value as described previously (Groen et al., 2020). For trait input data, we used the genotypic means of replicate measurements, which were square-root transformed prior to GWA mapping to ensure normality. For genotype input data, we used a previously described SNP dataset of 179,634 markers (Groen et al., 2020), which had been filtered so that SNP variants were excluded if they were inside regions overlapping repetitive regions or within 5 bp of an indel variant (raw FASTQ reads for all accessions are available from the NCBI’s Sequence Read Archive, https://www.ncbi.nlm.nih.gov/sra, under BioProject accession numbers PRJNA422249, PRJNA557122, and PRJEB6180). SNPs further needed to have at least 80% of accessions with a genotype call prior to missing genotype imputation, a minor allele frequency of at least 5%, and a heterozygous genotype in less than 5% of the accessions. Finally, the SNPs had been randomly pruned by sampling only a single polymorphic site within every 1,000-bp window. Within the MLMM, we included population structure cofactors as well as a kinship matrix as a random factor that had been constructed from the SNP dataset using the VanRaden method in GAPIT as described previously (Groen et al., 2020). We inferred population structure among the Indica accessions using principal component analysis (PCA) and let GAPIT use the first six PCs as cofactors, since sub-populations of Indica rice can start to be distinguished at PC6 (Fuentes et al., 2019). We obtained the significance of SNPs using a conservative Bonferroni threshold at p<2.78 × 10^−7^.

Identification of TGA TF-binding sites was accomplished by obtaining the genomic sequence of a 10kb region upstream and 5kb region downstream of the candidate genes *OsSEU1* and *OsSEU2* from the EnsemblPlants database (https://plants.ensembl.org/). For candidate gene *OsFbox536*, we selected a 25kb region that includes three tandemly duplicated F-box protein-encoding genes (*OsFbox535-537*) to search for TGA TF-binding sites. These sequences were read into Geneious Prime software, where we could search for and annotate occurrences of the TGA motif (TGACGT). Locations of motifs can be found with their genomic coordinates in Table S5.

### Mutant Analysis

A stable rice mutant line (CCRM-2208) for *OsTGA5*/*rTGA2.3* (Os01g0279900, LOC_Os01g17260) was generated by Creative Biogene (Shirley, NY) via CRISPR/Cas9 gene editing using binary vector pCB-35S-Cas9-OsU6-sgRNA-HygR in Japonica accession Nipponbare. Embryonic calli were transformed after plasmid transformation to *Agrobacterium tumefaciens* strain EHA105. The *tga5* loss-of-function mutation was confirmed with next-generation sequencing of a 2.2kb region using primers (F 5’GCTGCTTCTGACTCTGACAGA3’ and R 5’CTTTAGCCTGCTATTCTCCAATTGT3’) that encompass the gRNA target site (TCACTTTGTAGACATTGCGT) (Fig. S12). Linear/PCR sequencing was performed by Plasmidsaurus using Oxford Nanopore Technology sequencing with custom analysis and annotation. After seed bulking, seeds for the wildtype Nipponbare and *tga5* loss-of-function mutant (in the Nipponbare background) were stored at 5℃ and relative humidity <60% until use in experiments.

For drought experiments with the wildtype Nipponbare and *tga5* mutant, rice plants were cultivated in a greenhouse at the Plant Research 1 facility on the University of California Riverside campus at 28°C and relative humidity <60% under natural light conditions during the summer season as described previously (Dunivant et al., 2024). The plants were grown in 3.8-L pots containing 3 L of soil. The pots offered flooded soil conditions as in rice paddies, providing well-watered conditions. Intermittent drought treatment started when plants were two months old. For the intermittent drought, treatment plants were removed from their water source, allowing the soil to dry. Plants were left in droughted conditions until we noticed leaf rolling (on average after 3-5 days). Plants were then placed back into the well-watered conditions to recover (on average for 5-7 days). This treatment was repeated continuously until the end of panicle development, at approximately 3 months after starting the first drought treatment.

### Gene Expression Stochasticity Analyses

We previously assessed genome-wide gene expression variation in leaf blades of plants in the 4^th^ hills of the 660 plots in each of the wet and dry field environments at 33 DAT, 17 days after withholding water that led to drought conditions (Groen et al., 2020). Processed read counts (available at https://zenodo.org/records/3533431 with DOI 10.5281/zenodo.3533431) were scaled to transcript count per million, further processed via invariant-set normalization, and log_2_-transformed for assessment of genome-wide gene expression stochasticity (Groen et al., 2020). Only transcripts from protein-coding genes on nuclear chromosomes that were detected in at least 10% of individuals across our populations were considered, leaving a total of n=15,635 transcripts. We calculated estimates of each genotype’s global gene expression stochasticity per environment by calculating the squared coefficient of variation ( *CV^2^* = *σ^2^*/*μ^2^* ) for each transcript across the three replicate individuals of an accession in an environment and then averaging *CV^2^*across all transcripts (Groen et al., 2020).

We performed genotypic selection analysis on global gene expression stochasticity in each environment for the Indica population as described above for selection on trait canalization but this time without including genotypic mean expression as a covariate, since the *CV^2^* measure of expression stochasticity already takes this into account.

We determined genotypic correlations between global gene expression stochasticity and trait canalization as Pearson’s product-moment correlation coefficients using the cor function in R version 4.0.3.

### Source Environment Analysis

For associating trait canalization with climate parameters from accessions’ source environments, we extracted data on bioclimatic variables from the WorldClim database (https://www.worldclim.org). We determined genotypic correlations between climate parameters and trait canalization as Pearson’s product-moment correlation coefficients using the cor function in R version 4.0.3.

## Data, Materials, and Software Availability

DNA resequencing data (in the form of raw FASTQ reads) for all accessions are available from the NCBI’s Sequence Read Archive, https://www.ncbi.nlm.nih.gov/sra, under BioProject accession numbers PRJNA422249, PRJNA557122, and PRJEB6180. Processed RNA sequencing read counts for all accessions are available from Zenodo (https://zenodo.org/records/3533431 with DOI 10.5281/zenodo.3533431). Information on experimental design, accession metadata, and trait data from experiments described here can be found as supplemental datasets.

## Acknowledgements

We thank Mark Siegal, Elena Hamann, and members of the Henry, Purugganan, and Groen laboratories for helpful discussions. We thank Bianca Uzziel Principe, Paul Maturan, and Leo Holongbayan for assistance with field management, This research was supported by the US National Science Foundation with grant DGE-1922642 “NRT: Plants-3D (Discover, Design and Deploy): Enhancing Minority Graduate Training in Plant Synthetic Biology” providing a fellowship to T.S.D., an Agriculture and Food Research Initiative Predoctoral Fellowship from the US National Institute of Food and Agriculture (grant 1032584 to T.S.D.), the Gordon and Betty Moore Foundation (grant GBMF2550.06 to S.C.G.), the National Institute of General Medical Sciences of the National Institutes of Health (grant R35GM151194 to S.C.G.), University of California Riverside startup funds (to S.C.G.), the US National Science Foundation Plant Genome Research Program (grants IOS 1546218 and 2204374 to M.D.P.), the Zegar Family Foundation, and the Canada Research Chairs (grant CRC-2021-00126 to Z.J.L.).

## Supplemental Figure Legends

**Fig. S1.** Set up of the field experiment with 220 Indica and Japonica rice accessions and their source environments. (*A*) Experimental design of the field experiment with 220 rice accessions planted in triplicate in row plots in a well-watered, wet paddy and a rain-out shelter, in which plants were exposed to an intermittent drought stress, according to an identical alpha-lattice design in each environment. Each plot consisted of a single 2-m row of 10 isogenic plants and plants were spaced 0.2 m apart from any neighboring plant. (*B*) Geographic distributions of source environments of all rice accessions included in the field experiment. Indica accessions are indicated in purple and Japonica accessions in green.

**Fig. S2.** Drought impacts trait means in Indica rice accessions. Frequency distributions for genotypic means of 11 morphological traits in Indica accessions (n=93). Blue indicates wet conditions and orange indicates dry conditions. From a two-way ANOVA, environment was significant for all traits (p<0.05).

**Fig. S3.** Drought induces trait decanalization in Indica rice accessions. Frequency distributions for trait canalization (as measured through micro-environmental variation) for six morphological traits in Indica accessions (n=93). Blue indicates wet conditions and orange indicates dry conditions. From a two-way ANOVA, environment was significant for all traits (p<0.05).

**Fig. S4.** Drought impacts trait means in Japonica rice accessions. Frequency distributions for genotypic means of 11 morphological traits in Japonica accessions (n=50). Blue indicates wet conditions and orange indicates dry conditions. From a two-way ANOVA, environment was significant for all traits (p<0.05).

**Fig. S5.** Drought induces trait decanalization in Japonica rice accessions. Frequency distributions for trait canalization (as measured through micro-environmental variation) for 11 morphological traits in Japonica accessions (n=50). Blue indicates wet conditions and orange indicates dry conditions. From a two-way ANOVA, environment was significant for all traits (p<0.05), except for tiller number, for which the environmental effect was marginally non-significant (p=0.06345).

**Fig. S6.** Genotypic correlations between trait means and canalization (as measured through micro-environmental variation) for six morphological traits in Indica rice accessions. Blue indicates wet conditions and orange indicates dry conditions. No significant Pearson product-moment correlations were identified (p>0.05).

**Fig. S7.** Genotypic correlations between trait means and canalization (as measured through micro-environmental variation) for 11 morphological traits in Japonica rice accessions. Blue indicates wet conditions and orange indicates dry conditions. No significant Pearson product-moment correlations were identified (p>0.05).

**Fig. S8.** Reproductive-stage traits quantified for wild-type Nipponbare and *tga5* loss-of-function mutant plants exposed to a 3-month period of intermittent drought or a well-watered control environment. Final measurements of trait means for plant height (*A*), tiller number (*B*), and shoot dry weight (*C*). Images of plants at the end of flowering for wild-type plants in wet (*D*) and dry (*E*) conditions and *tga5* mutant plants in wet (*F*) and dry (*G*) conditions. Blue indicates wet conditions and orange indicates dry conditions. From a two-way ANOVA, genotype and environment were significant for plant height: p=7.36×10-6, p=0.0127; tiller number: p=1.27×10-6, p=0.00608; and shoot dry weight: p=0.000182, p=0.000395, respectively.

**Fig. S9.** Panicle traits quantified for wild-type Nipponbare and *tga5* loss-of-function mutant plants exposed to a 3-month period of intermittent drought or a well-watered control environment. (*A*-*B*) Final measurements of trait means and canalization (as measured through micro-environmental variation) for spikelet number per panicle. Statistically significant factors (genotype, environment or genotype-by-environment interaction) from a two-way ANOVA are indicated below each panel. (*C*) Images of representative panicles and individual panicle branches at the end of flowering for wild-type and *tga5* mutant plants in wet (blue) and dry (orange) conditions.

**Fig. S10.** Panicle traits quantified for wild-type Nipponbare and *tga5* loss-of-function mutant plants exposed to a 3-month period of intermittent drought or a well-watered control environment. (*A*-*D*) Final measurements of trait means and canalization (as measured through micro-environmental variation) for panicles’ primary (*A*-*B*) and secondary branch numbers (*C*-*D*), respectively. Statistically significant factors (genotype, environment or genotype-by-environment interaction) from a two-way ANOVA are indicated below each panel. Blue indicates wet conditions and orange indicates dry conditions.

**Fig. S11.** Panicle traits quantified for wild-type Nipponbare and *tga5* loss-of-function mutant plants exposed to a 3-month period of intermittent drought or a well-watered control environment. (*A*-*D*) Final measurements of trait means and canalization (as measured through micro-environmental variation) for panicles’ percentages of unfilled grains (*A*-*B*). Statistically significant factors (genotype, environment or genotype-by-environment interaction) from a two-way ANOVA are indicated below each panel and images of representative panicles are shown. Blue indicates wet conditions and orange indicates dry conditions.

**Fig. S12.** Rice *tga5* loss-of-function mutant plants contain a single base pair insertion resulting in a premature stop codon in exon 4. (*A*) Gene model of *OsTGA5* annotated with the TGA-binding domain DOG1 in orange, and gRNA, mutation site, and stop codon annotated with blue arrows. The 5’ and 3’ UTR are indicated in grey, exons in black, and introns with a dashed line. (*B*) DNA and AA sequences of the Nipponbare reference and the *tga5* mutant with the single-bp insertion highlighted by a dotted box. (*C*) DNA and AA sequence of the Nipponbare reference with the predicted single-bp frameshift in exon 4 where we find a premature stop codon indicated with an arrow.

## Supplemental Tables

**Table S1.** Within-environment broad-sense heritabilities (*H^2^*) for trait means and canalization in the Indica population.

**Table S2.** Broad-sense heritabilities (*H^2^*) for trait means and canalization in the Japonica population.

**Table S3.** Selection gradients (*β*) for trait means and canalization in the Indica population.

**Table S4.** Selection gradients (*β*) for trait means and canalization in the Japonica population.

**Table S5.** TGA motif coordinates for the three candidate loci as annotated on EnsemblPlants.

## Supplemental Data

**Data S1.** Metadata for Indica and Japonica rice accessions.

**Data S2.** Raw trait data, the trait median and log-transformed Levene’s Statistic (LS, canalization) values per plot for 11 morphological traits, and fecundity fitness values per plot of all Indica and Japonica accessions in the wet field environment.

**Data S3.** Raw trait data, the trait median and log-transformed Levene’s Statistic (LS, canalization) values per plot for 11 morphological traits, and fecundity fitness values per plot of all Indica and Japonica accessions in the dry field environment.

**Data S4.** Data on the trait median and log-transformed Levene’s Statistic (LS, canalization) values per plot for 11 morphological traits and on fecundity fitness of Indica and Japonica accessions in the wet and dry field environments used for data analysis.

**Data S5.** Genome-wide association analyses on genotypic mean values of the trait median and log-transformed Levene’s Statistic (LS, canalization) values per plot for 11 morphological traits of Indica accessions in the wet and dry field environments.

**Data S6.** Data on the trait mean and log-transformed Levene’s Statistic (LS, canalization) values for wild-type Nipponbare and *tga5* loss-of-function mutant plants exposed to a 3-month period of intermittent drought or a well-watered control environment.

**Data S7.** Genotypic selection analyses on global gene expression stochasticity in leaves and correlation analyses between genotypic mean values of the trait median and log-transformed Levene’s Statistic (LS, canalization) values per plot for 11 morphological traits and leaf expression stochasticity as well as source environment characteristics of Indica accessions.

## References

1. V. Debat, P. David, Mapping phenotypes: canalization, plasticity and developmental stability. Trends Ecol Evol 16, 555–561 (2001).

2. C. D. Schlichting, M. Pigliucci. Phenotypic evolution. A reaction norm perspective. Sunderland, UK: Sinauer Associates Inc (1998).

3. J. D. Glover, et al., The developmental basis of fingerprint pattern formation and variation. Cell 186, 940–956.e20 (2023).

4. B. G. Forde, Is it good noise? The role of developmental instability in the shaping of a root system. J Exp Bot 60, 3989–4002 (2009).

5. J. Lachowiec, C. Queitsch, D. J. Kliebenstein, Molecular mechanisms governing differential robustness of development and environmental responses in plants. Ann Bot 117, 795–809 (2016).

6. M. Viney, S. E. Reece, Adaptive noise. Proc Biol Sci 280, 20131104 (2013).

7. L. Mestek Boukhibar, M. Barkoulas, The developmental genetics of biological robustness. Ann Bot 117, 699–707 (2016).

8. T. Flatt, The evolutionary genetics of canalization. Q Rev Biol 80, 287–316 (2005).

9. B. Hallgrimsson, et al., The developmental genetics of canalization. Semin Cell Dev Biol 88, 67–79 (2019).

10. M. Bruijning, C. J. E. Metcalf, E. Jongejans, J. F. Ayroles, The evolution of variance control. Trends Ecol Evol 35, 22–33 (2020).

11. S. A. Diaz, M. Viney, Genotypic-specific variance in *Caenorhabditis elegans* lifetime fecundity. Ecol Evol 4, 2058–2069 (2014).

12. J. F. Ayroles, et al., Behavioral idiosyncrasy reveals genetic control of phenotypic variability. Proc Natl Acad Sci U S A 112, 6706–6711 (2015).

13. J. Chen, V. Nolte, C. Schlötterer, Temperature stress mediates decanalization and dominance of gene expression in *Drosophila melanogaster*. PLoS Genet 11, e1004883 (2015).

14. B. Chen, A. Wagner, Hsp90 is important for fecundity, longevity, and buffering of cryptic deleterious variation in wild fly populations. BMC Evol Biol 12, 25 (2012).

15. I. Dworkin, A study of canalization and developmental stability in the sternopleural bristle system of *Drosophila melanogaster*. Evolution 59, 1500–1509 (2005).

16. J. B. Lack, M. J. Monette, E. J. Johanning, Q. D. Sprengelmeyer, J. E. Pool, Decanalization of wing development accompanied the evolution of large wings in high-altitude *Drosophila*. Proc Natl Acad Sci U S A 113, 1014–1019 (2016).

17. F. Morgante, P. Sørensen, D. A. Sorensen, C. Maltecca, T. F. C. Mackay, Genetic architecture of micro-environmental plasticity in *Drosophila melanogaster*. Sci Rep 5, 9785 (2015).

18. J. B. Wang, H.-L. Lu, R. J. St Leger, The genetic basis for variation in resistance to infection in the *Drosophila melanogaster* genetic reference panel. PLoS Pathog 13, e1006260 (2017).

19. M. C. Hall, I. Dworkin, M. C. Ungerer, M. Purugganan, Genetics of microenvironmental canalization in *Arabidopsis thaliana*. Proc Natl Acad Sci U S A 104, 13717–13722 (2007).

20. J. M. Jimenez-Gomez, J. A. Corwin, B. Joseph, J. N. Maloof, D. J. Kliebenstein, Genomic analysis of QTLs and genes altering natural variation in stochastic noise. PLoS Genet 7, e1002295 (2011).

21. B. Joseph, J. A. Corwin, D. J. Kliebenstein, Genetic variation in the nuclear and organellar genomes modulates stochastic variation in the metabolome, growth, and defense. PLoS Genet 11, e1004779 (2015).

22. M. Kiskowski, T. Glimm, N. Moreno, T. Gamble, Y. Chiari, Isolating and quantifying the role of developmental noise in generating phenotypic variation. PLoS Comput Biol 15, e1006943 (2019).

23. H. A. Mulder, P. Gienapp, M. E. Visser, Genetic variation in variability: phenotypic variability of fledging weight and its evolution in a songbird population. Evolution 70, 2004–2016 (2016).

24. S. L. Rutherford, S. Lindquist, Hsp90 as a capacitor for morphological evolution. Nature 396, 336–342 (1998).

25. C. Queitsch, T. A. Sangster, S. Lindquist, Hsp90 as a capacitor of phenotypic variation. Nature 417, 618–624 (2002).

26. C. C. Milton, B. Huynh, P. Batterham, S. L. Rutherford, A. A. Hoffmann, Quantitative trait symmetry independent of Hsp90 buffering: distinct modes of genetic canalization and developmental stability. Proc Natl Acad Sci U S A 100, 13396–13401 (2003).

27. S. Alseekh, et al., Canalization of tomato fruit metabolism. Plant Cell 29, 2753–2765 (2017).

28. J. Fisher, E. Bensal, D. Zamir, Bimodality of stable and plastic traits in plants. Theor Appl Genet 130, 1915–1926 (2017).

29. H. Li, et al., Genetic variants and underlying mechanisms influencing variance heterogeneity in maize. Plant J 103, 1089–1102 (2020).

30. M. Sobral, J. Guitián, P. Guitián, A. R. Larrinaga, Seed predators exert selection on the subindividual variation of seed size. Plant Biol (Stuttg) 16, 836–842 (2014).

31. M. Sobral, J. Guitián, P. Guitián, C. Violle, A. R. Larrinaga, Exploring sub-individual variability: role of ontogeny, abiotic environment and seed-dispersing birds. Plant Biol (Stuttg) 21, 688–694 (2019).

32. C. Pélabon, et al., Is there more to within-plant variation in seed size than developmental noise? Evol Biol 48,366–377 (2021).

33. A. Beral, R. Rincent, J. Le Gouis, C. Girousse, V. Allard, Wheat individual grain-size variance originates from crop development and from specific genetic determinism. PLoS One 15, e0230689 (2020).

34. S. C. Groen, et al., The strength and pattern of natural selection on gene expression in rice. Nature 578, 572–576 (2020).

35. I. Ćalić, et al., The influence of genetic architecture on responses to selection under drought in rice. Evol Appl 15, 1670–1690 (2022).

36. E. Hamann, et al., Selection on genome-wide gene expression plasticity of rice in wet and dry field environments. Mol Ecol, e17522 (2024).

37. R. Lande, S. J. Arnold, The measurement of selection on correlated characters. Evolution 37, 1210–1226 (1983).

38. M. D. Rausher, The measurement of selection on quantitative traits: biases due to environmental covariances between traits and fitness. Evolution 46, 616–626 (1992).

39. A. Krishnan, C. Gupta, M. M. R. Ambavaram, A. Pereira, RECoN: Rice Environment Coexpression Network for systems level analysis of abiotic-stress response. Front Plant Sci 8, 1640 (2017).

40. C. Lekklar, et al., Genome-wide association study for salinity tolerance at the flowering stage in a panel of rice accessions from Thailand. BMC Genomics 20, 76 (2019).

41. W. Tanaka, T. Toriba, H.-Y. Hirano, Three *TOB1*-related *YABBY* genes are required to maintain proper function of the spikelet and branch meristems in rice. New Phytol 215, 825–839 (2017).

42. Y. Niu, et al., Phosphorylation of OsTGA5 by casein kinase II compromises its suppression of defense-related gene transcription in rice. Plant Cell 34, 3425–3442 (2022).

43. Y. Sato, et al., RiceFREND: a platform for retrieving coexpressed gene networks in rice. Nucleic Acids Res 41, D1214–21 (2013).

44. S. Ballouz, et al., The transcriptional legacy of developmental stochasticity. Nat Commun 14, 7226 (2023).

45. S. Cortijo, Z. Aydin, S. Ahnert, J. C. Locke, Widespread inter-individual gene expression variability in *Arabidopsis thaliana*. Mol Syst Biol 15, e8591 (2019).

46. S. Gupta, et al., Systems genomics of salinity stress response in rice. Elife 13 (2025).

47. A. Lea, et al., Genetic and environmental perturbations lead to regulatory decoherence. Elife 8 (2019).

48. M. B. Cruzan, M.A. Streisfeld, J.A. Schwoch, Fitness effects of somatic mutations accumulating during vegetative growth. Evol Ecol 36, 767–785 (2022).

49. C. Goeldel, F. Johannes, Stochasticity in gene body methylation. Curr Opin Plant Biol 75, 102436 (2023).

50. L. Wang, et al., The architecture of intra-organism mutation rate variation in plants. PLoS Biol 17, e3000191 (2019).

51. C. Yang, et al., OsLUGL is involved in the regulating auxin level and OsARFs expression in rice (*Oryza sativa* L.). Plant Sci 288, 110239 (2019).

52. X. Yin, et al., OsMADS18, a membrane-bound MADS-box transcription factor, modulates plant architecture and the abscisic acid response in rice. J Exp Bot 70, 3895–3909 (2019).

53. S. R. Yadav, I. Khanday, B. B. Majhi, K. Veluthambi, U. Vijayraghavan, Auxin-responsive OsMGH3, a common downstream target of OsMADS1 and OsMADS6, controls rice floret fertility. Plant Cell Physiol 52, 2123–2135 (2011).

54. G. Xu, et al., uORF-mediated translation allows engineered plant disease resistance without fitness costs. Nature 545, 491–494 (2017).

55. X. Li, et al., The systemic acquired resistance regulator OsNPR1 attenuates growth by repressing auxin signaling through promoting IAA-amido synthase expression. Plant Physiol 172, 546–558 (2016).

56. B. Wang, et al., Condensation of SEUSS promotes hyperosmotic stress tolerance in *Arabidopsis*. Nat Chem Biol 18, 1361–1369 (2022).

57. N. P. R. Anten, P. J. Vermeulen, Tragedies and crops: understanding natural selection to improve cropping systems. Trends Ecol Evol 31, 429–439 (2016).

58. R. M. Gutaker, et al., Genomic history and ecology of the geographic spread of rice. Nat Plants 6, 492–502 (2020).

59. R. R. Fuentes, et al., Structural variants in 3000 rice genomes. Genome Res 29, 870–880 (2019).

60. T. S. Dunivant, et al., Evolutionary systems biology identifies genetic trade-offs in rice defense against above- and belowground attackers. Plant Cell Physiol (2024). 10.1093/pcp/pcae107.

