## Supplemental Figures and Tables for "Environment-dependent selection impacts heritable developmental stability and trait canalization in rice"

### Slide 1
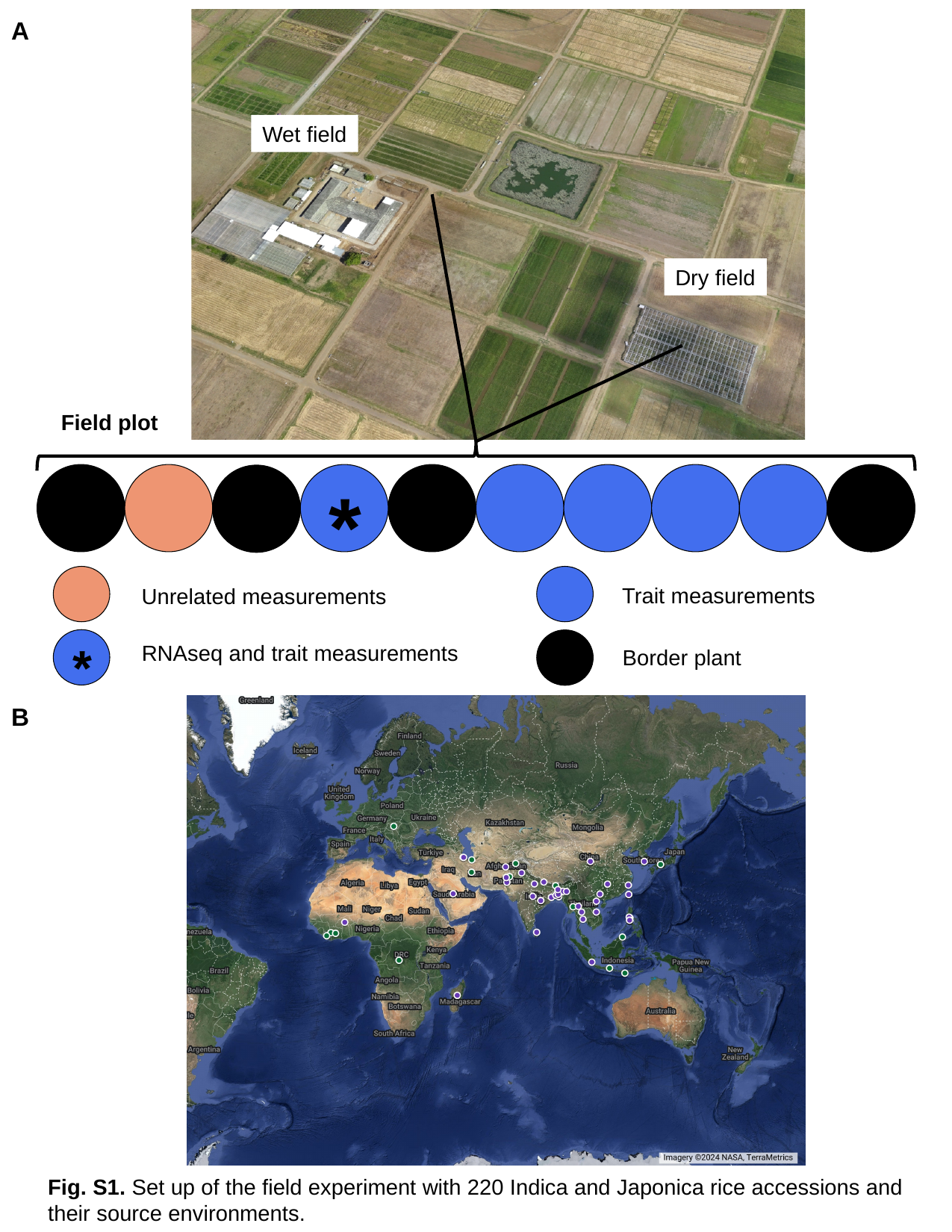

A
Wet field
Dry field
Field plot
*
Trait measurements
Unrelated measurements
*
RNAseq and trait measurements
Border plant
B
Fig. S1. Set up of the field experiment with 220 Indica and Japonica rice accessions and their source environments.

### Slide 2
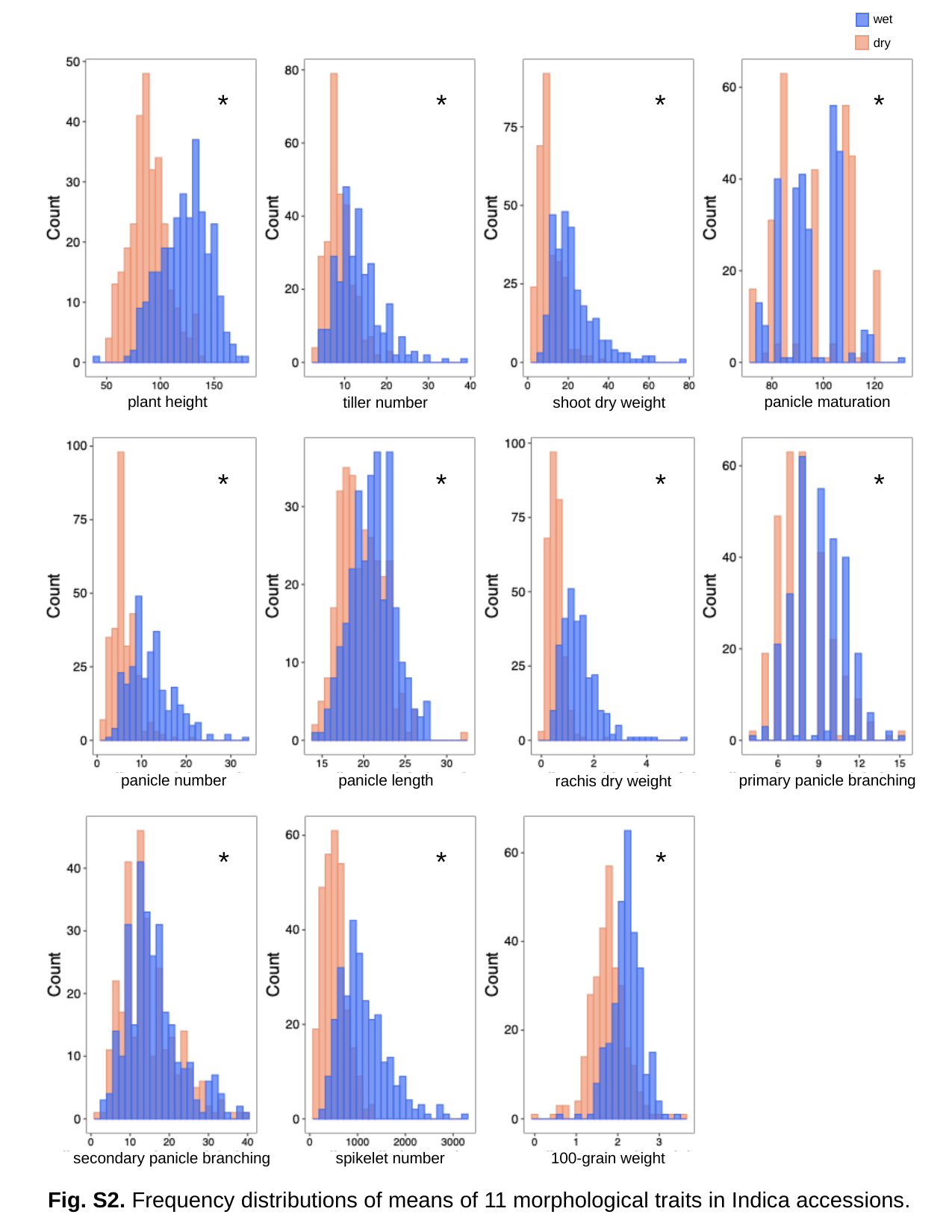

wet
dry
plant height
panicle maturation
shoot dry weight
tiller number
panicle number
primary panicle branching
panicle length
rachis dry weight
100-grain weight
secondary panicle branching
spikelet number
*
*
*
*
*
*
*
*
*
*
*
Fig. S2. Frequency distributions of means of 11 morphological traits in Indica accessions.

### Slide 3
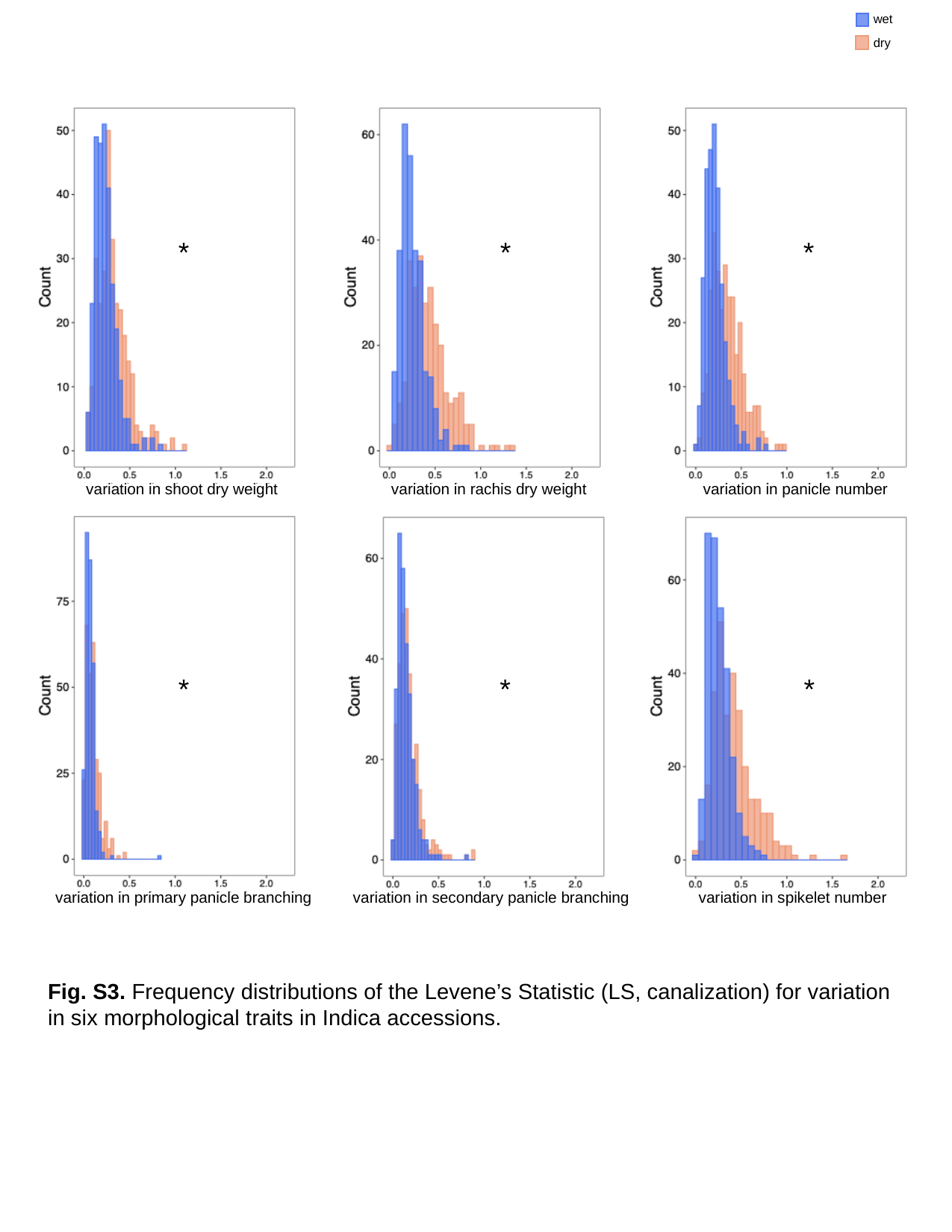

wet
dry
variation in shoot dry weight
variation in rachis dry weight
variation in panicle number
variation in primary panicle branching
variation in secondary panicle branching
variation in spikelet number
*
*
*
*
*
*
Fig. S3. Frequency distributions of the Levene’s Statistic (LS, canalization) for variation in six morphological traits in Indica accessions.

### Slide 4
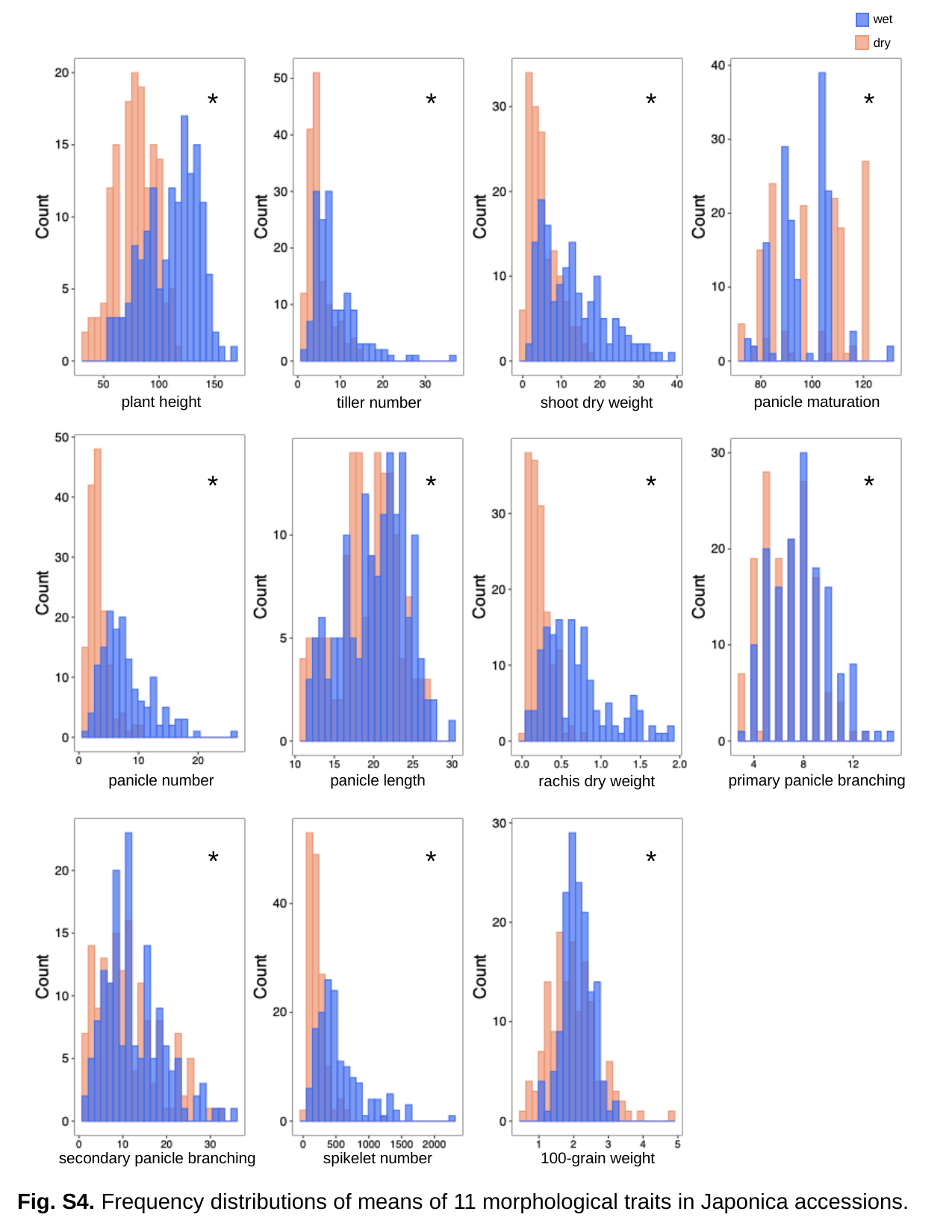

wet
dry
plant height
panicle maturation
shoot dry weight
tiller number
panicle number
primary panicle branching
panicle length
rachis dry weight
100-grain weight
secondary panicle branching
spikelet number
*
*
*
*
*
*
*
*
*
*
*
Fig. S4. Frequency distributions of means of 11 morphological traits in Japonica accessions.

### Slide 5
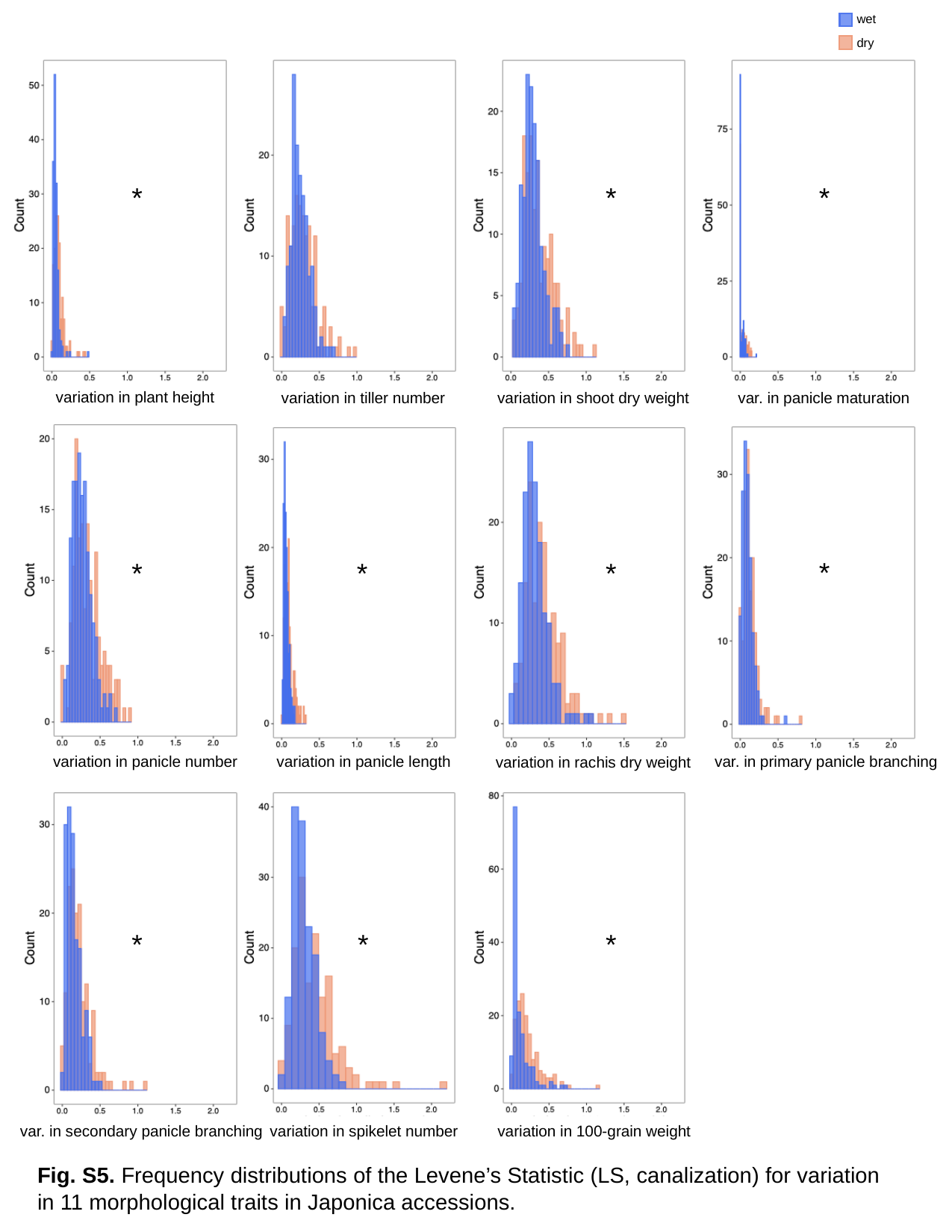

wet
dry
variation in plant height
var. in panicle maturation
variation in shoot dry weight
variation in tiller number
variation in panicle number
var. in primary panicle branching
variation in panicle length
variation in rachis dry weight
variation in 100-grain weight
var. in secondary panicle branching
variation in spikelet number
*
*
*
*
*
*
*
*
*
*
Fig. S5. Frequency distributions of the Levene’s Statistic (LS, canalization) for variation in 11 morphological traits in Japonica accessions.

### Slide 6
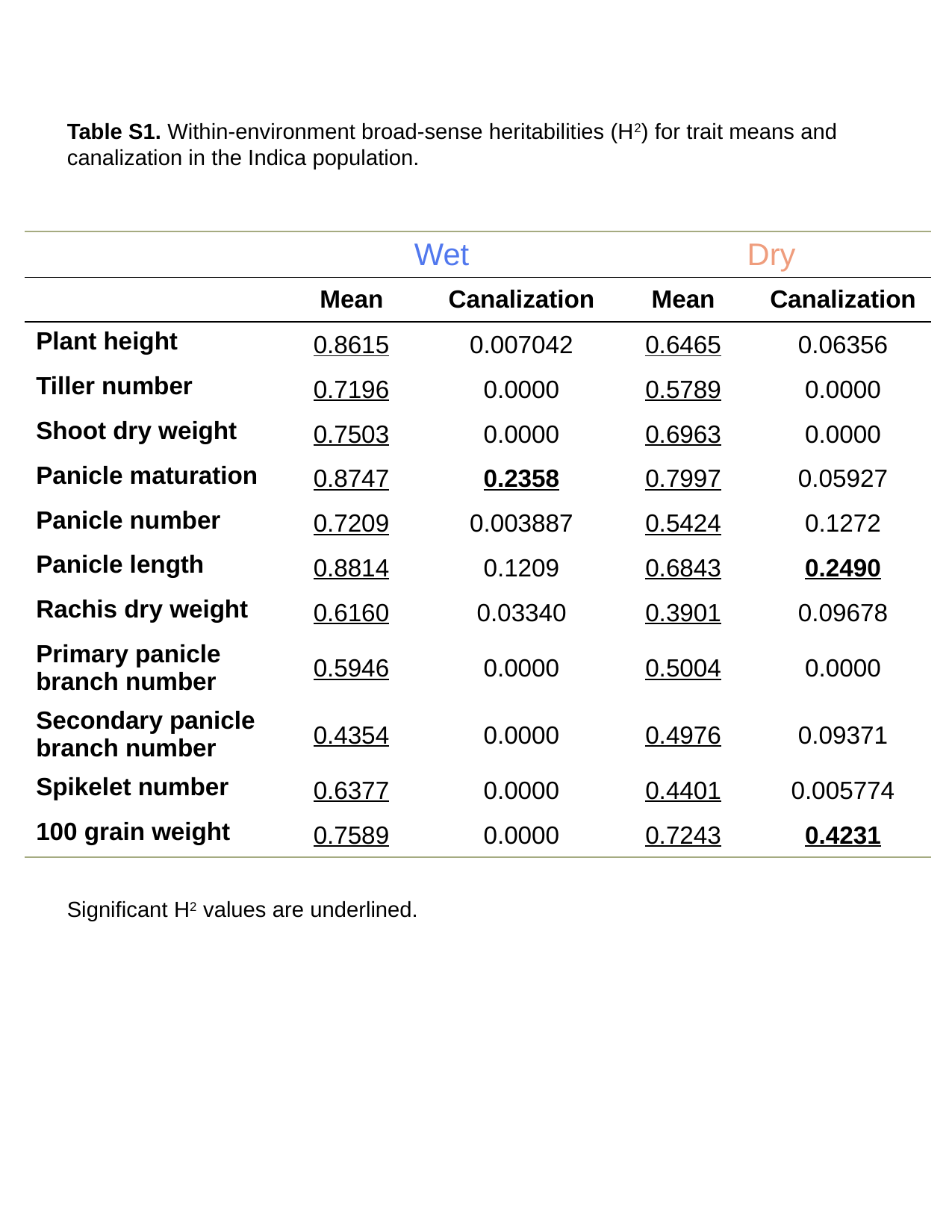

Table S1. Within-environment broad-sense heritabilities (H^2^) for trait means and canalization in the Indica population.
| | Wet | | Dry | Dry |
| --- | --- | --- | --- | --- |
| | Mean | Canalization | Mean | Canalization |
| Plant height | 0.8615 | 0.007042 | 0.6465 | 0.06356 |
| Tiller number | 0.7196 | 0.0000 | 0.5789 | 0.0000 |
| Shoot dry weight | 0.7503 | 0.0000 | 0.6963 | 0.0000 |
| Panicle maturation | 0.8747 | 0.2358 | 0.7997 | 0.05927 |
| Panicle number | 0.7209 | 0.003887 | 0.5424 | 0.1272 |
| Panicle length | 0.8814 | 0.1209 | 0.6843 | 0.2490 |
| Rachis dry weight | 0.6160 | 0.03340 | 0.3901 | 0.09678 |
| Primary panicle branch number | 0.5946 | 0.0000 | 0.5004 | 0.0000 |
| Secondary panicle branch number | 0.4354 | 0.0000 | 0.4976 | 0.09371 |
| Spikelet number | 0.6377 | 0.0000 | 0.4401 | 0.005774 |
| 100 grain weight | 0.7589 | 0.0000 | 0.7243 | 0.4231 |
Significant H^2^ values are underlined.

### Slide 7
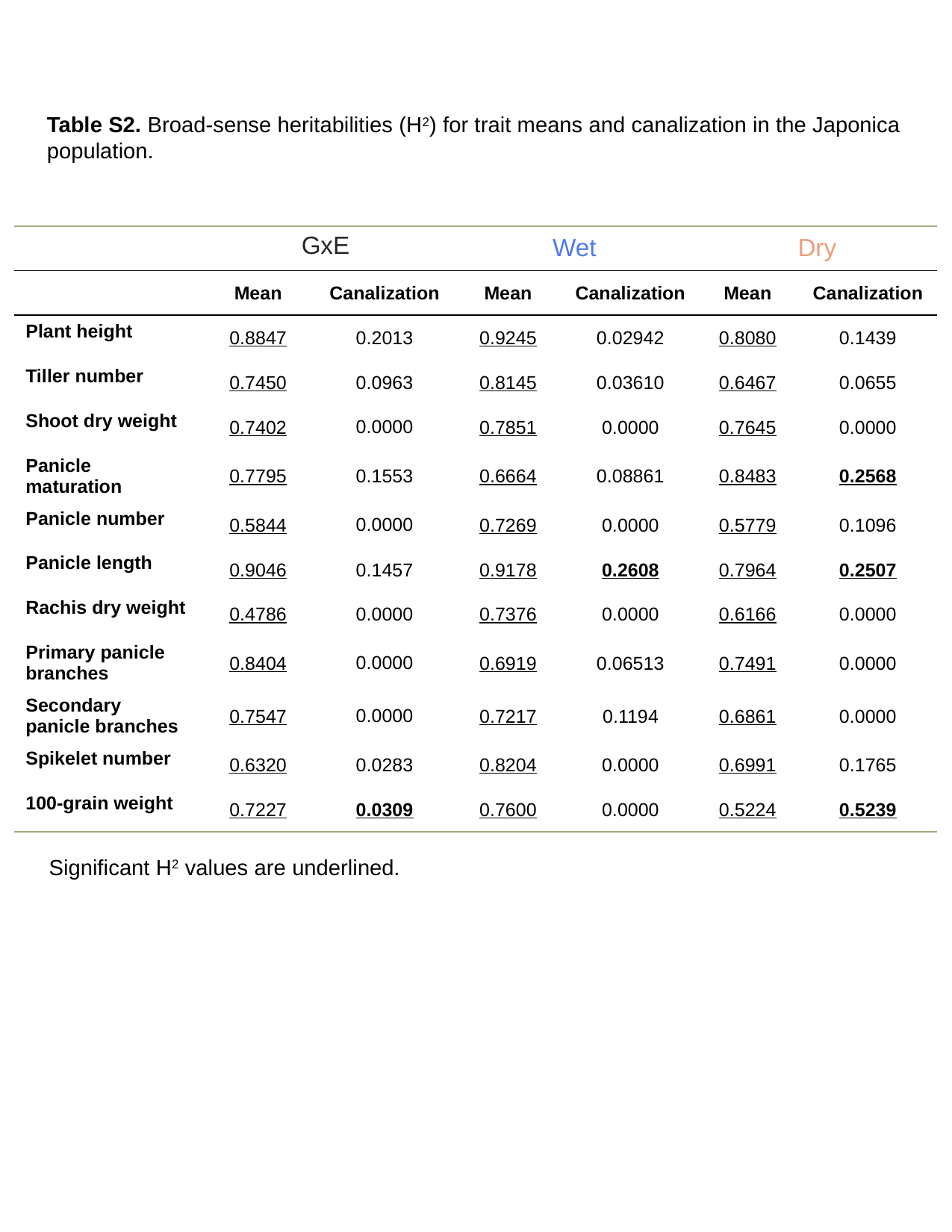

Table S2. Broad-sense heritabilities (H^2^) for trait means and canalization in the Japonica population.
| | GxE | | Wet | | Dry | Dry |
| --- | --- | --- | --- | --- | --- | --- |
| | Mean | Canalization | Mean | Canalization | Mean | Canalization |
| Plant height | 0.8847 | 0.2013 | 0.9245 | 0.02942 | 0.8080 | 0.1439 |
| Tiller number | 0.7450 | 0.0963 | 0.8145 | 0.03610 | 0.6467 | 0.0655 |
| Shoot dry weight | 0.7402 | 0.0000 | 0.7851 | 0.0000 | 0.7645 | 0.0000 |
| Panicle maturation | 0.7795 | 0.1553 | 0.6664 | 0.08861 | 0.8483 | 0.2568 |
| Panicle number | 0.5844 | 0.0000 | 0.7269 | 0.0000 | 0.5779 | 0.1096 |
| Panicle length | 0.9046 | 0.1457 | 0.9178 | 0.2608 | 0.7964 | 0.2507 |
| Rachis dry weight | 0.4786 | 0.0000 | 0.7376 | 0.0000 | 0.6166 | 0.0000 |
| Primary panicle branches | 0.8404 | 0.0000 | 0.6919 | 0.06513 | 0.7491 | 0.0000 |
| Secondary panicle branches | 0.7547 | 0.0000 | 0.7217 | 0.1194 | 0.6861 | 0.0000 |
| Spikelet number | 0.6320 | 0.0283 | 0.8204 | 0.0000 | 0.6991 | 0.1765 |
| 100-grain weight | 0.7227 | 0.0309 | 0.7600 | 0.0000 | 0.5224 | 0.5239 |
Significant H^2^ values are underlined.

### Slide 8
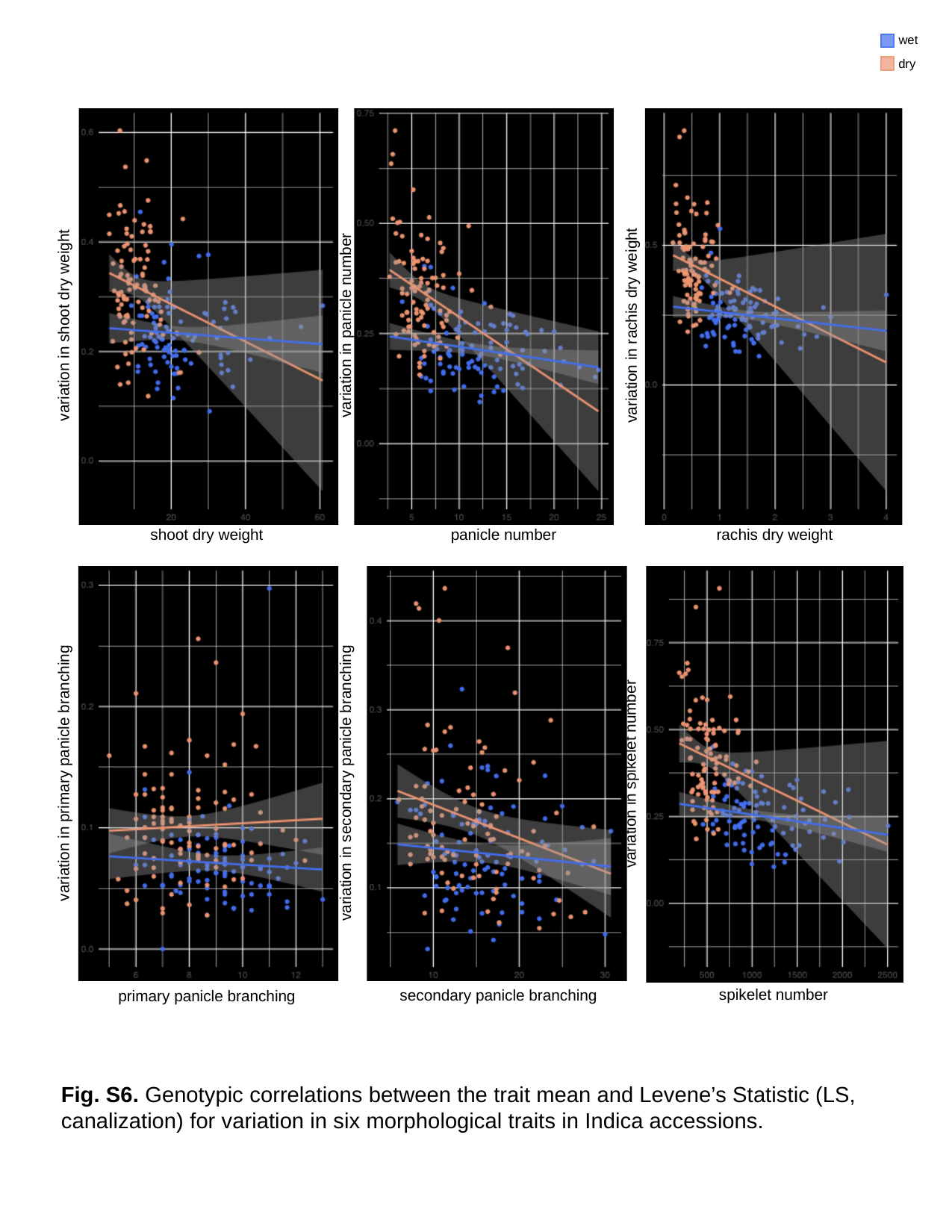

wet
dry
variation in shoot dry weight
variation in rachis dry weight
variation in panicle number
panicle number
rachis dry weight
shoot dry weight
variation in primary panicle branching
variation in spikelet number
variation in secondary panicle branching
spikelet number
secondary panicle branching
primary panicle branching
Fig. S6. Genotypic correlations between the trait mean and Levene’s Statistic (LS, canalization) for variation in six morphological traits in Indica accessions.

### Slide 9
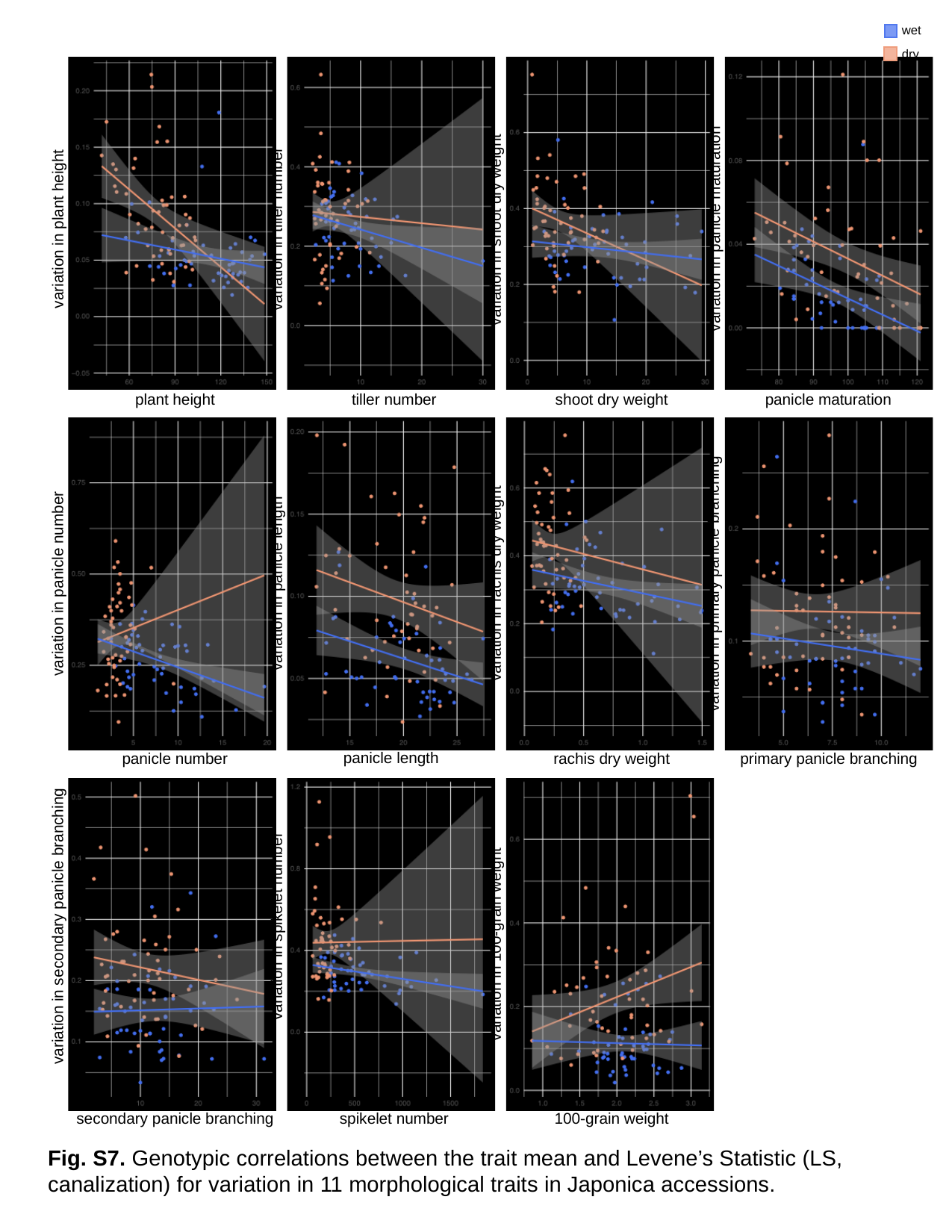

wet
dry
variation in panicle maturation
variation in tiller number
variation in plant height
variation in shoot dry weight
panicle maturation
plant height
tiller number
shoot dry weight
variation in rachis dry weight
variation in panicle length
variation in panicle number
variation in primary panicle branching
panicle length
primary panicle branching
panicle number
rachis dry weight
variation in spikelet number
variation in secondary panicle branching
variation in 100-grain weight
secondary panicle branching
spikelet number
100-grain weight
Fig. S7. Genotypic correlations between the trait mean and Levene’s Statistic (LS, canalization) for variation in 11 morphological traits in Japonica accessions.

### Slide 10
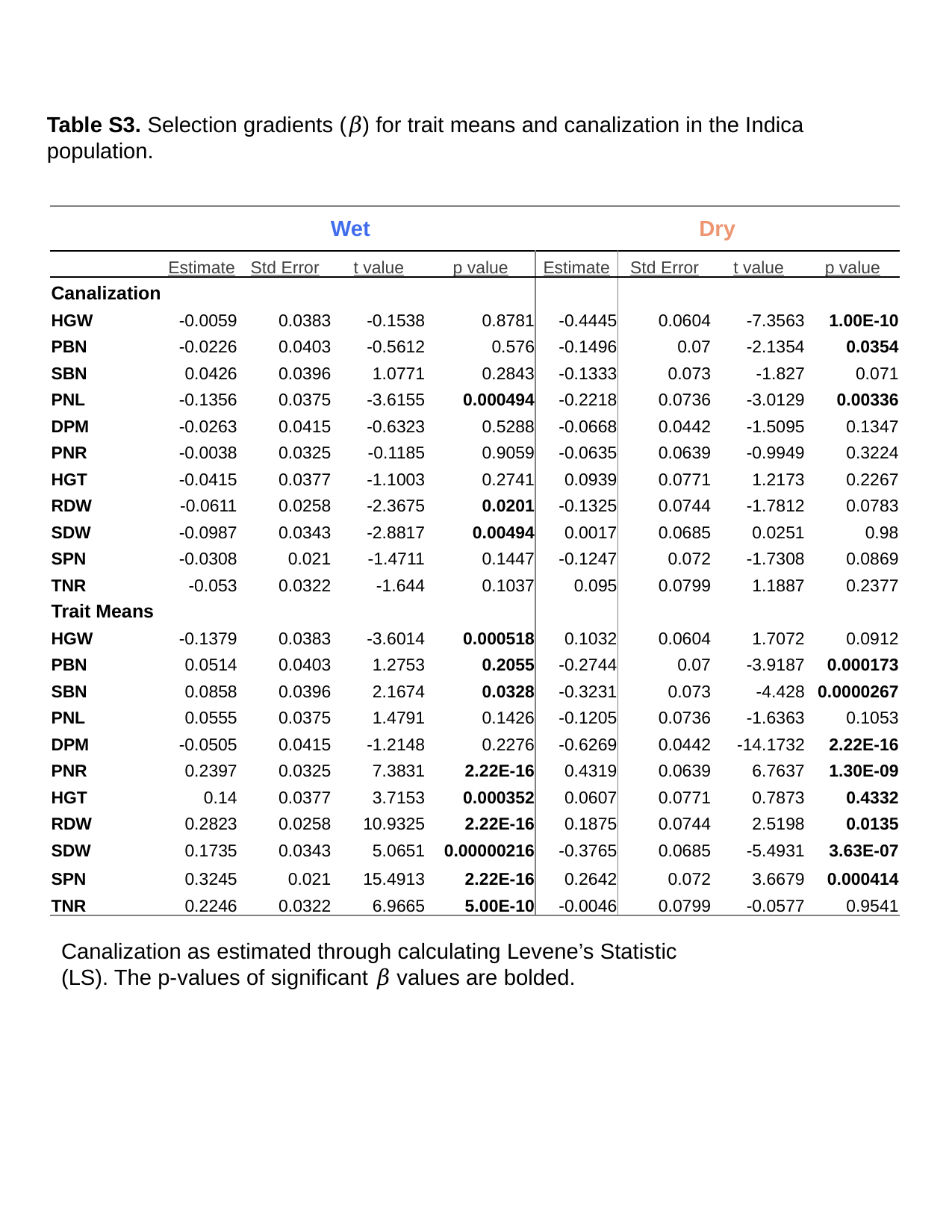

Table S3. Selection gradients (𝛽) for trait means and canalization in the Indica population.
| | Wet | | | | Dry | | | |
| --- | --- | --- | --- | --- | --- | --- | --- | --- |
| | Estimate | Std Error | t value | p value | Estimate | Std Error | t value | p value |
| Canalization | | | | | | | | |
| HGW | -0.0059 | 0.0383 | -0.1538 | 0.8781 | -0.4445 | 0.0604 | -7.3563 | 1.00E-10 |
| PBN | -0.0226 | 0.0403 | -0.5612 | 0.576 | -0.1496 | 0.07 | -2.1354 | 0.0354 |
| SBN | 0.0426 | 0.0396 | 1.0771 | 0.2843 | -0.1333 | 0.073 | -1.827 | 0.071 |
| PNL | -0.1356 | 0.0375 | -3.6155 | 0.000494 | -0.2218 | 0.0736 | -3.0129 | 0.00336 |
| DPM | -0.0263 | 0.0415 | -0.6323 | 0.5288 | -0.0668 | 0.0442 | -1.5095 | 0.1347 |
| PNR | -0.0038 | 0.0325 | -0.1185 | 0.9059 | -0.0635 | 0.0639 | -0.9949 | 0.3224 |
| HGT | -0.0415 | 0.0377 | -1.1003 | 0.2741 | 0.0939 | 0.0771 | 1.2173 | 0.2267 |
| RDW | -0.0611 | 0.0258 | -2.3675 | 0.0201 | -0.1325 | 0.0744 | -1.7812 | 0.0783 |
| SDW | -0.0987 | 0.0343 | -2.8817 | 0.00494 | 0.0017 | 0.0685 | 0.0251 | 0.98 |
| SPN | -0.0308 | 0.021 | -1.4711 | 0.1447 | -0.1247 | 0.072 | -1.7308 | 0.0869 |
| TNR | -0.053 | 0.0322 | -1.644 | 0.1037 | 0.095 | 0.0799 | 1.1887 | 0.2377 |
| Trait Means | | | | | | | | |
| HGW | -0.1379 | 0.0383 | -3.6014 | 0.000518 | 0.1032 | 0.0604 | 1.7072 | 0.0912 |
| PBN | 0.0514 | 0.0403 | 1.2753 | 0.2055 | -0.2744 | 0.07 | -3.9187 | 0.000173 |
| SBN | 0.0858 | 0.0396 | 2.1674 | 0.0328 | -0.3231 | 0.073 | -4.428 | 0.0000267 |
| PNL | 0.0555 | 0.0375 | 1.4791 | 0.1426 | -0.1205 | 0.0736 | -1.6363 | 0.1053 |
| DPM | -0.0505 | 0.0415 | -1.2148 | 0.2276 | -0.6269 | 0.0442 | -14.1732 | 2.22E-16 |
| PNR | 0.2397 | 0.0325 | 7.3831 | 2.22E-16 | 0.4319 | 0.0639 | 6.7637 | 1.30E-09 |
| HGT | 0.14 | 0.0377 | 3.7153 | 0.000352 | 0.0607 | 0.0771 | 0.7873 | 0.4332 |
| RDW | 0.2823 | 0.0258 | 10.9325 | 2.22E-16 | 0.1875 | 0.0744 | 2.5198 | 0.0135 |
| SDW | 0.1735 | 0.0343 | 5.0651 | 0.00000216 | -0.3765 | 0.0685 | -5.4931 | 3.63E-07 |
| SPN | 0.3245 | 0.021 | 15.4913 | 2.22E-16 | 0.2642 | 0.072 | 3.6679 | 0.000414 |
| TNR | 0.2246 | 0.0322 | 6.9665 | 5.00E-10 | -0.0046 | 0.0799 | -0.0577 | 0.9541 |
Canalization as estimated through calculating Levene’s Statistic (LS). The p-values of significant 𝛽 values are bolded.

### Slide 11
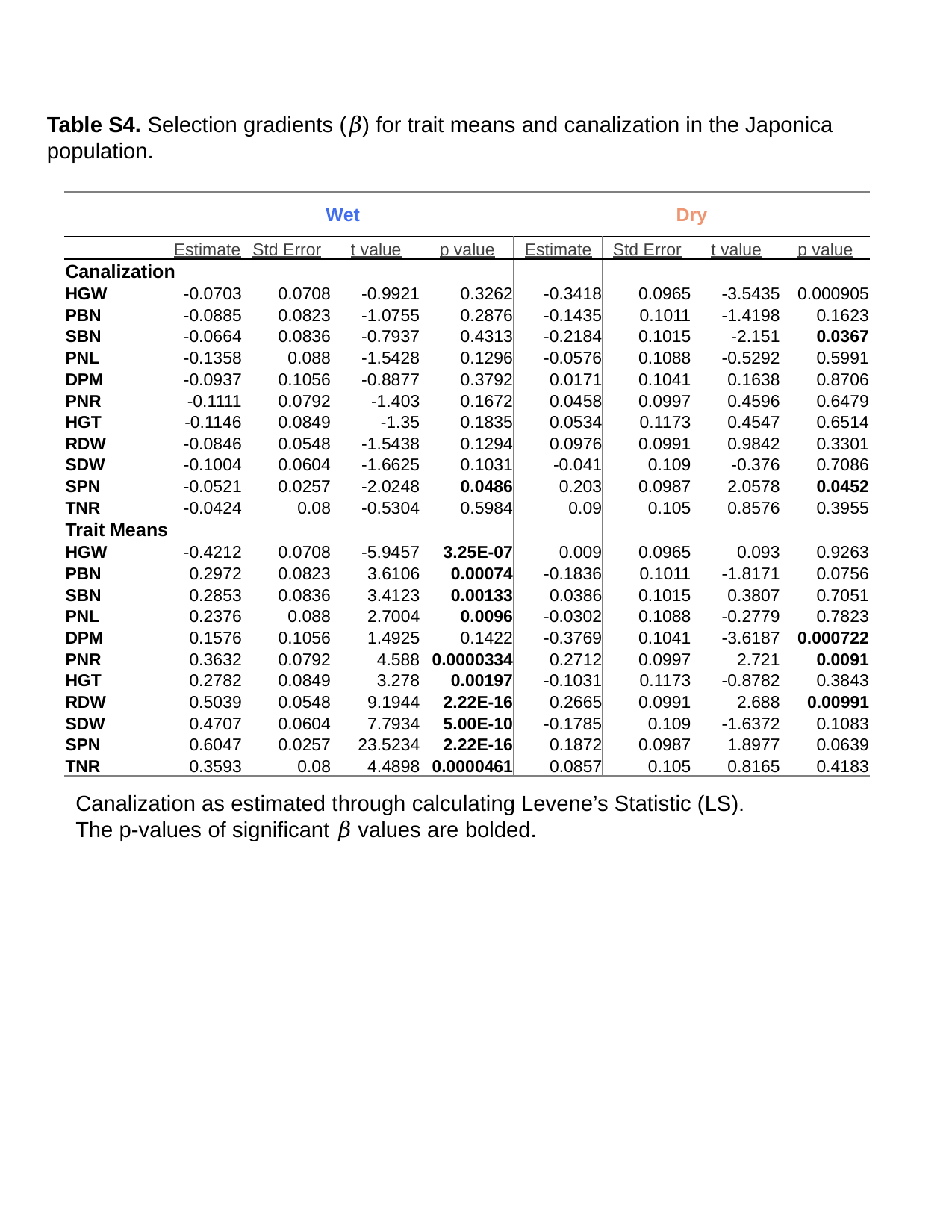

Table S4. Selection gradients (𝛽) for trait means and canalization in the Japonica population.
| | Wet | | | | Dry | | | |
| --- | --- | --- | --- | --- | --- | --- | --- | --- |
| | Estimate | Std Error | t value | p value | Estimate | Std Error | t value | p value |
| Canalization | | | | | | | | |
| HGW | -0.0703 | 0.0708 | -0.9921 | 0.3262 | -0.3418 | 0.0965 | -3.5435 | 0.000905 |
| PBN | -0.0885 | 0.0823 | -1.0755 | 0.2876 | -0.1435 | 0.1011 | -1.4198 | 0.1623 |
| SBN | -0.0664 | 0.0836 | -0.7937 | 0.4313 | -0.2184 | 0.1015 | -2.151 | 0.0367 |
| PNL | -0.1358 | 0.088 | -1.5428 | 0.1296 | -0.0576 | 0.1088 | -0.5292 | 0.5991 |
| DPM | -0.0937 | 0.1056 | -0.8877 | 0.3792 | 0.0171 | 0.1041 | 0.1638 | 0.8706 |
| PNR | -0.1111 | 0.0792 | -1.403 | 0.1672 | 0.0458 | 0.0997 | 0.4596 | 0.6479 |
| HGT | -0.1146 | 0.0849 | -1.35 | 0.1835 | 0.0534 | 0.1173 | 0.4547 | 0.6514 |
| RDW | -0.0846 | 0.0548 | -1.5438 | 0.1294 | 0.0976 | 0.0991 | 0.9842 | 0.3301 |
| SDW | -0.1004 | 0.0604 | -1.6625 | 0.1031 | -0.041 | 0.109 | -0.376 | 0.7086 |
| SPN | -0.0521 | 0.0257 | -2.0248 | 0.0486 | 0.203 | 0.0987 | 2.0578 | 0.0452 |
| TNR | -0.0424 | 0.08 | -0.5304 | 0.5984 | 0.09 | 0.105 | 0.8576 | 0.3955 |
| Trait Means | | | | | | | | |
| HGW | -0.4212 | 0.0708 | -5.9457 | 3.25E-07 | 0.009 | 0.0965 | 0.093 | 0.9263 |
| PBN | 0.2972 | 0.0823 | 3.6106 | 0.00074 | -0.1836 | 0.1011 | -1.8171 | 0.0756 |
| SBN | 0.2853 | 0.0836 | 3.4123 | 0.00133 | 0.0386 | 0.1015 | 0.3807 | 0.7051 |
| PNL | 0.2376 | 0.088 | 2.7004 | 0.0096 | -0.0302 | 0.1088 | -0.2779 | 0.7823 |
| DPM | 0.1576 | 0.1056 | 1.4925 | 0.1422 | -0.3769 | 0.1041 | -3.6187 | 0.000722 |
| PNR | 0.3632 | 0.0792 | 4.588 | 0.0000334 | 0.2712 | 0.0997 | 2.721 | 0.0091 |
| HGT | 0.2782 | 0.0849 | 3.278 | 0.00197 | -0.1031 | 0.1173 | -0.8782 | 0.3843 |
| RDW | 0.5039 | 0.0548 | 9.1944 | 2.22E-16 | 0.2665 | 0.0991 | 2.688 | 0.00991 |
| SDW | 0.4707 | 0.0604 | 7.7934 | 5.00E-10 | -0.1785 | 0.109 | -1.6372 | 0.1083 |
| SPN | 0.6047 | 0.0257 | 23.5234 | 2.22E-16 | 0.1872 | 0.0987 | 1.8977 | 0.0639 |
| TNR | 0.3593 | 0.08 | 4.4898 | 0.0000461 | 0.0857 | 0.105 | 0.8165 | 0.4183 |
Canalization as estimated through calculating Levene’s Statistic (LS). The p-values of significant 𝛽 values are bolded.

### Slide 12
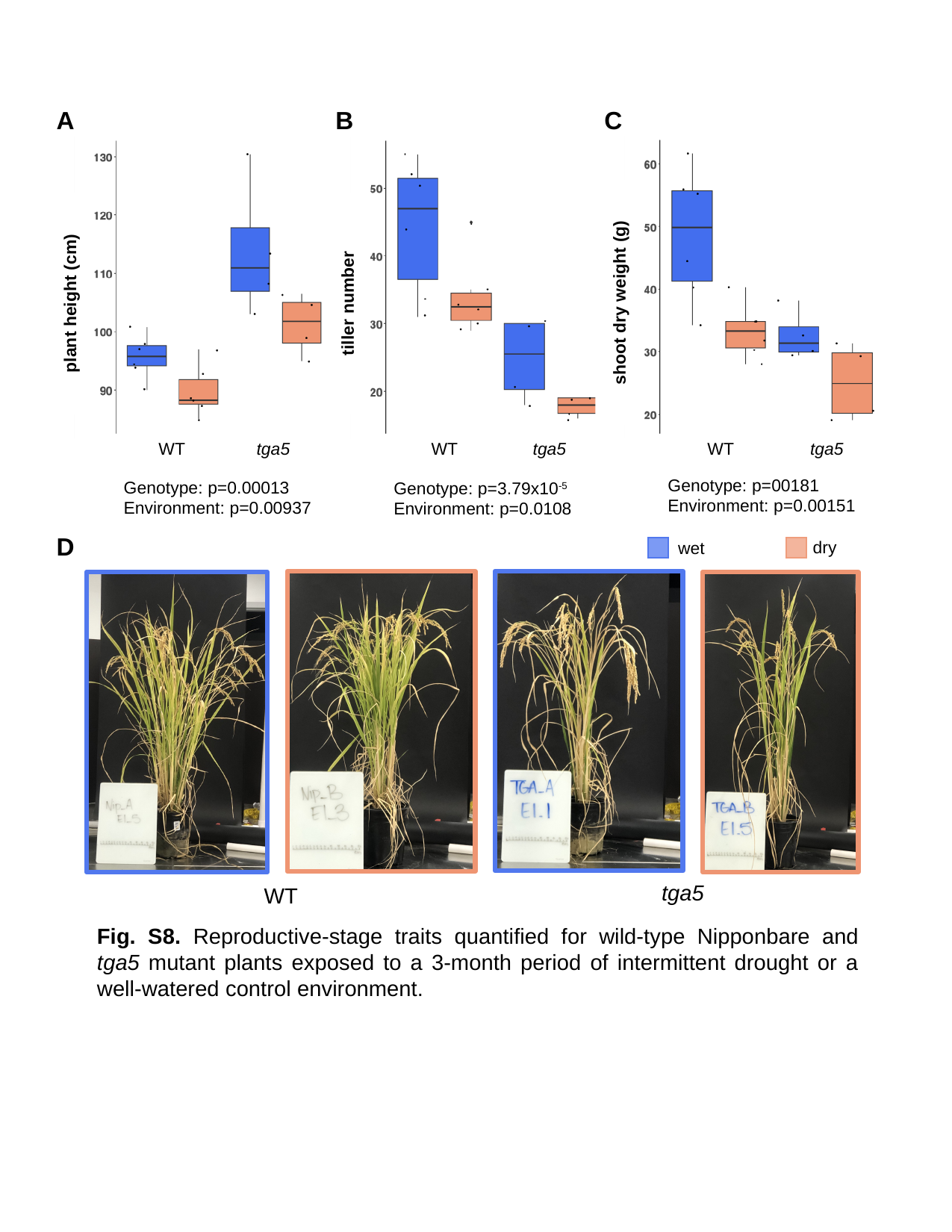

A
B
C
shoot dry weight (g)
plant height (cm)
tiller number
tga5
WT
tga5
WT
tga5
WT
D
dry
wet
tga5
WT
Genotype: p=00181
Environment: p=0.00151
Genotype: p=0.00013
Environment: p=0.00937
Genotype: p=3.79x10^-5^
Environment: p=0.0108
Fig. S8. Reproductive-stage traits quantified for wild-type Nipponbare and tga5 mutant plants exposed to a 3-month period of intermittent drought or a well-watered control environment.

### Slide 13
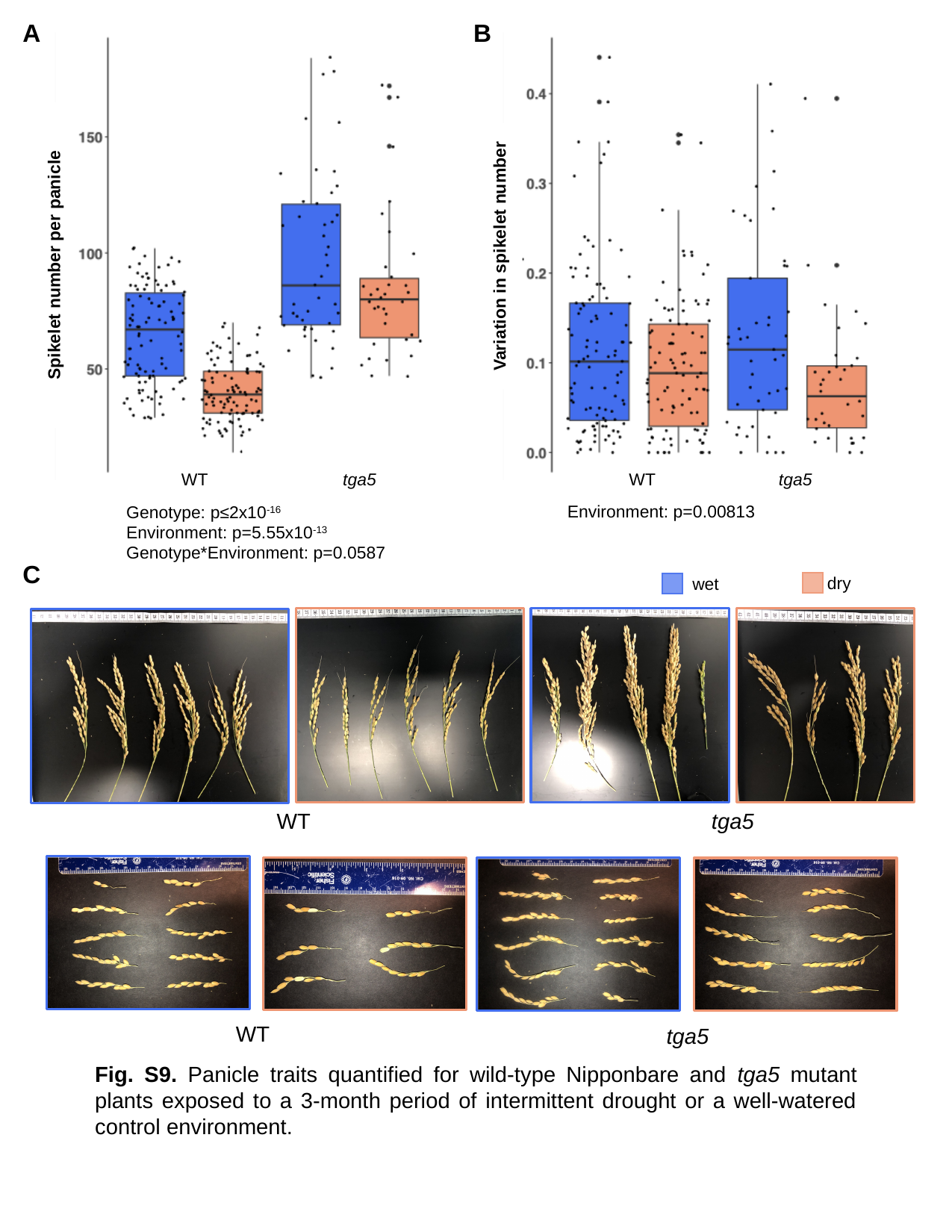

A
B
WT
tga5
WT
tga5
Environment: p=0.00813
Genotype: p≤2x10^-16^
Environment: p=5.55x10^-13^
Genotype*Environment: p=0.0587
C
dry
wet
WT
tga5
WT
tga5
Variation in spikelet number
Spikelet number per panicle
Fig. S9. Panicle traits quantified for wild-type Nipponbare and tga5 mutant plants exposed to a 3-month period of intermittent drought or a well-watered control environment.

### Slide 14
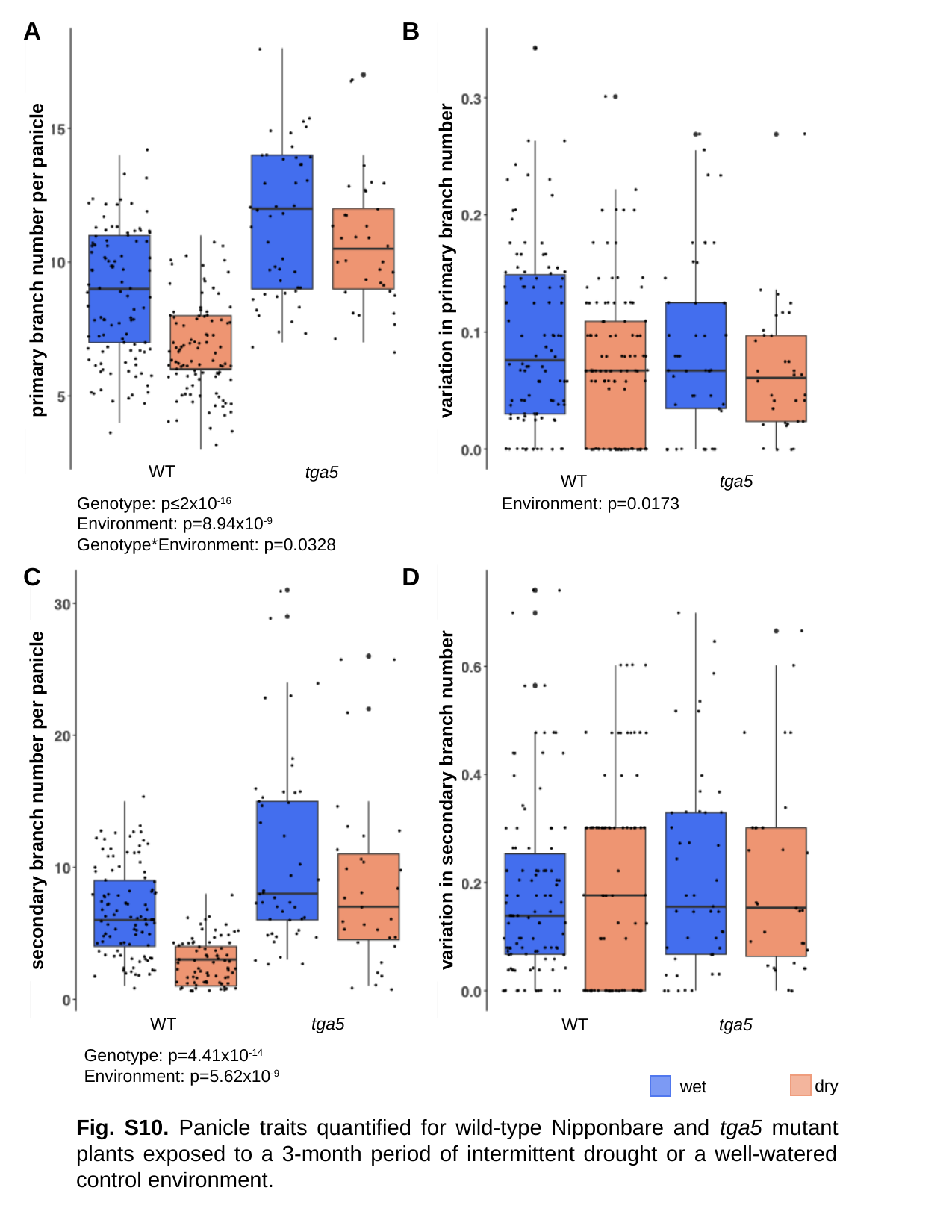

A
B
WT
tga5
tga5
WT
Environment: p=0.0173
C
D
variation in secondary branch number
tga5
WT
tga5
WT
Genotype: p=4.41x10^-14^
Environment: p=5.62x10^-9^
dry
wet
primary branch number per panicle
variation in primary branch number
Genotype: p≤2x10^-16^
Environment: p=8.94x10^-9^
Genotype*Environment: p=0.0328
secondary branch number per panicle
Fig. S10. Panicle traits quantified for wild-type Nipponbare and tga5 mutant plants exposed to a 3-month period of intermittent drought or a well-watered control environment.

### Slide 15
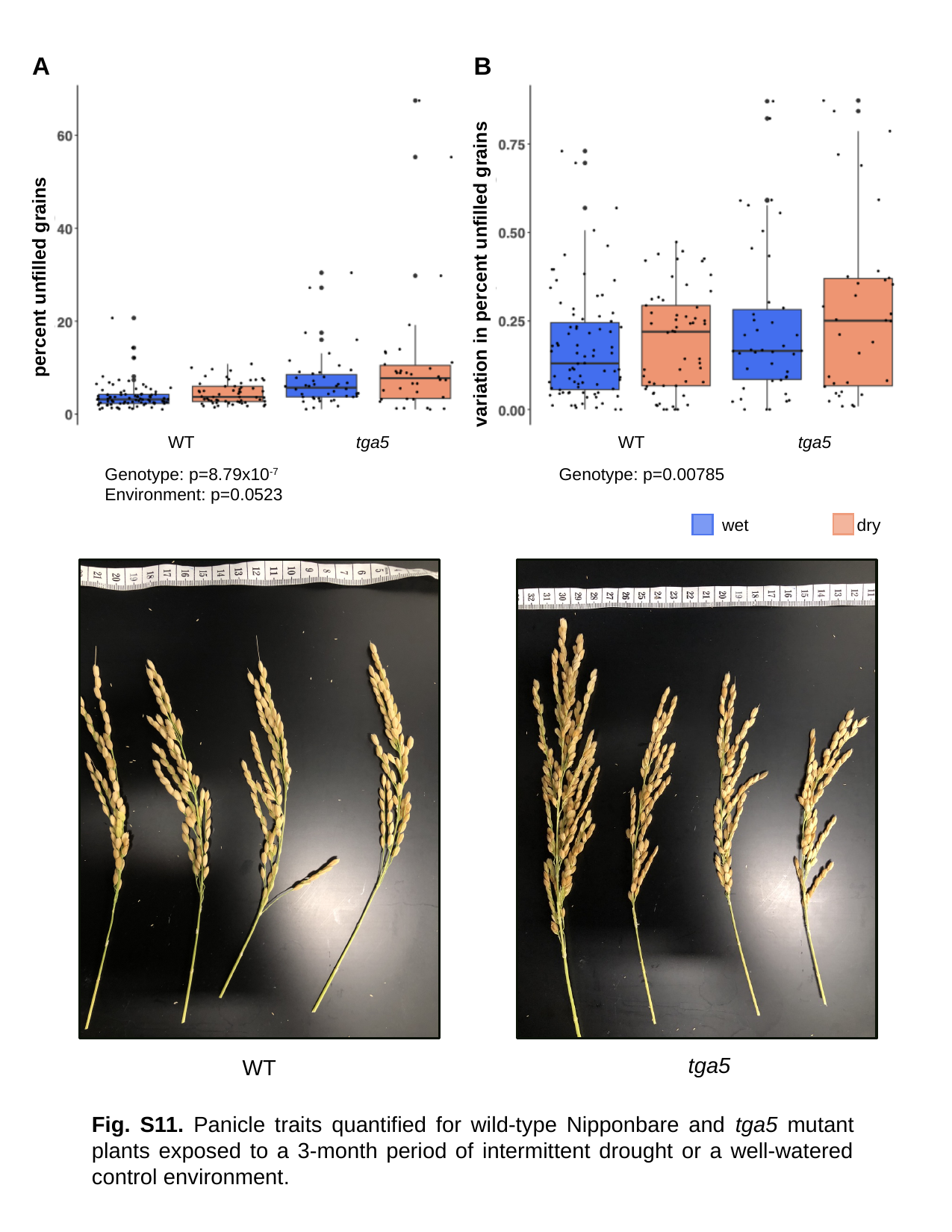

A
B
variation in percent unfilled grains
percent unfilled grains
WT
tga5
WT
tga5
Genotype: p=8.79x10^-7^
Environment: p=0.0523
Genotype: p=0.00785
dry
wet
tga5
WT
Fig. S11. Panicle traits quantified for wild-type Nipponbare and tga5 mutant plants exposed to a 3-month period of intermittent drought or a well-watered control environment.

### Slide 16
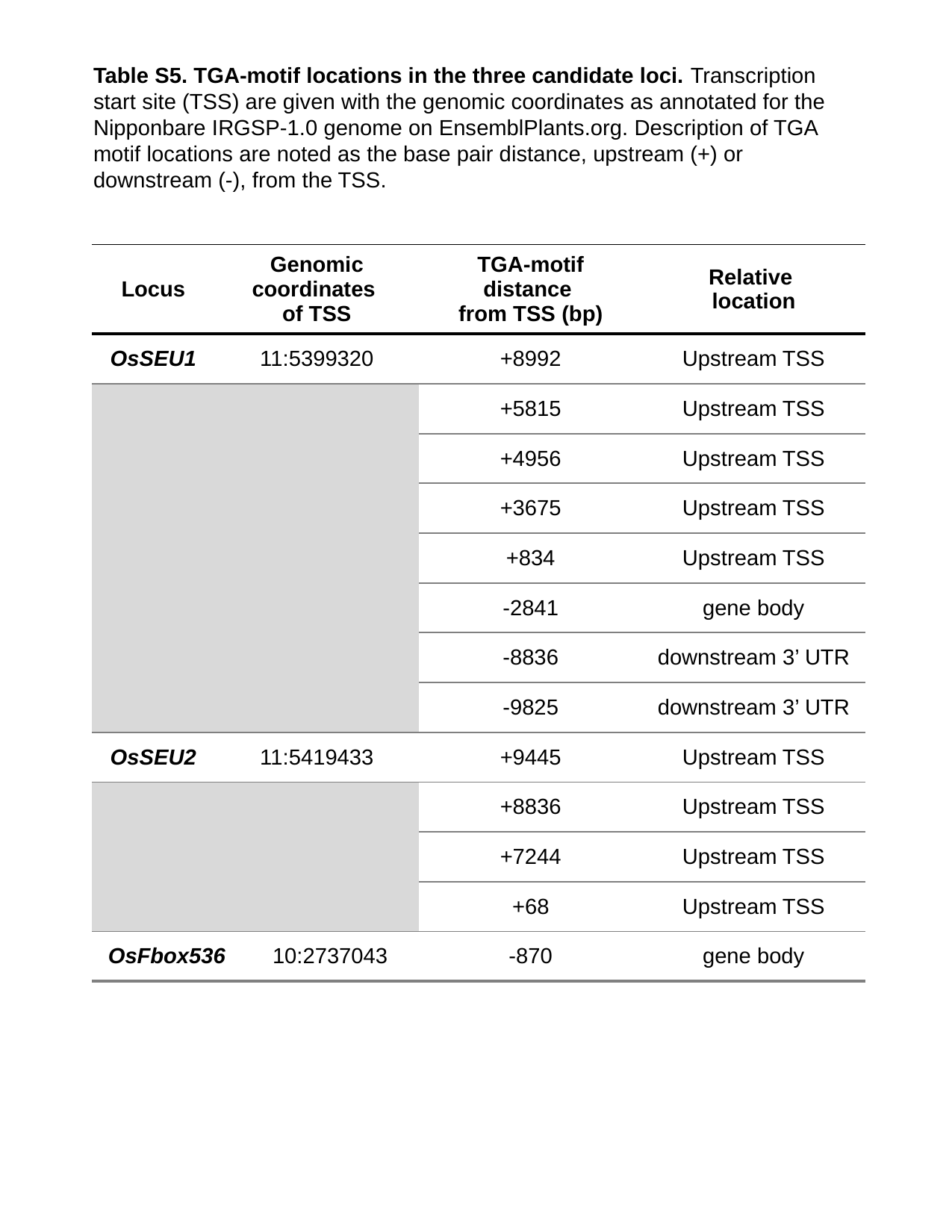

Table S5. TGA-motif locations in the three candidate loci. Transcription start site (TSS) are given with the genomic coordinates as annotated for the Nipponbare IRGSP-1.0 genome on EnsemblPlants.org. Description of TGA motif locations are noted as the base pair distance, upstream (+) or downstream (-), from the TSS.
| Locus | Genomic coordinates of TSS | | TGA-motif distance from TSS (bp) | Relative location |
| --- | --- | --- | --- | --- |
| OsSEU1 | 11:5399320 | | +8992 | Upstream TSS |
| | | | +5815 | Upstream TSS |
| | | | +4956 | Upstream TSS |
| | | | +3675 | Upstream TSS |
| | | | +834 | Upstream TSS |
| | | | -2841 | gene body |
| | | | -8836 | downstream 3’ UTR |
| | | | -9825 | downstream 3’ UTR |
| OsSEU2 | 11:5419433 | | +9445 | Upstream TSS |
| | | | +8836 | Upstream TSS |
| | | | +7244 | Upstream TSS |
| | | | +68 | Upstream TSS |
| OsFbox536 | 10:2737043 | 10:2737043 | -870 | gene body |

### Slide 17
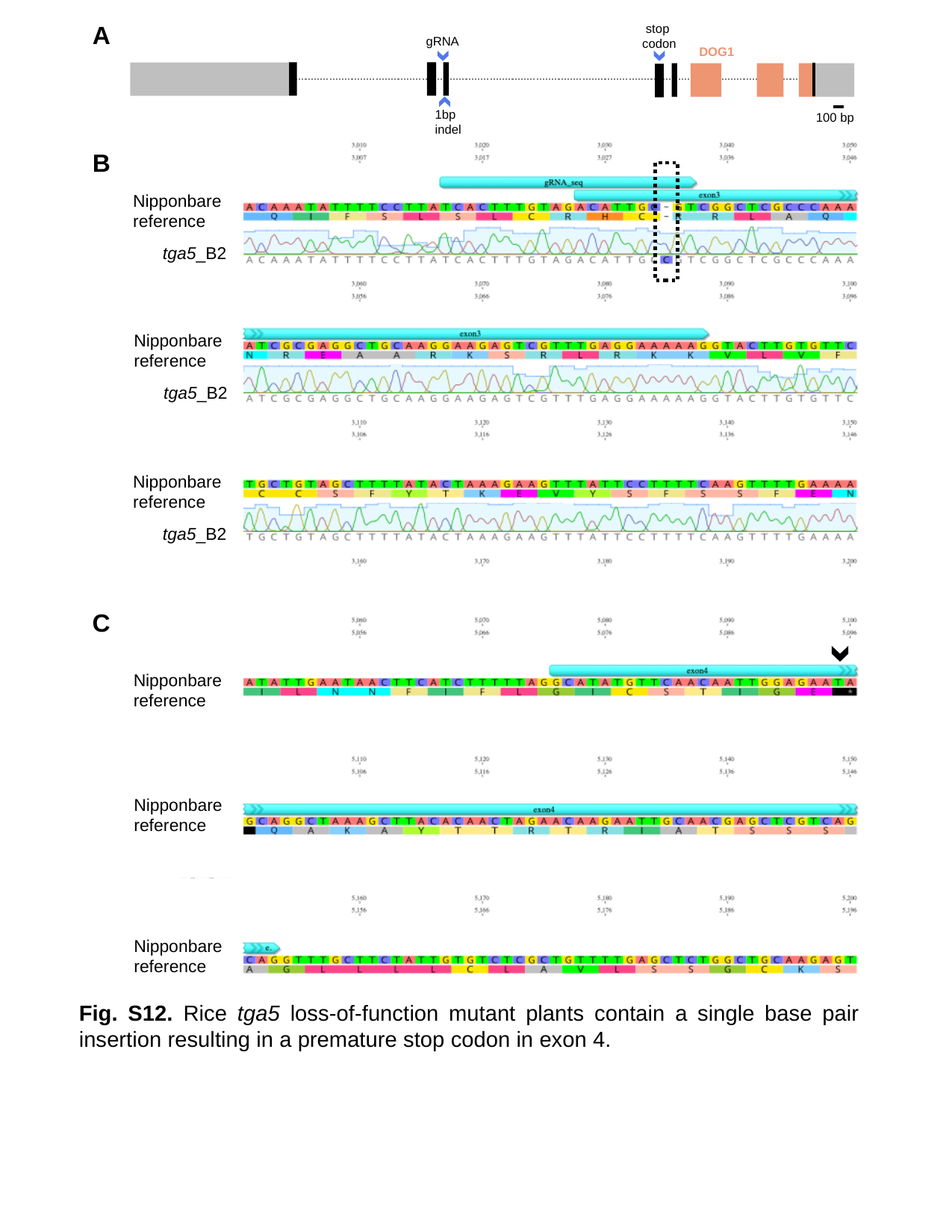

A
stop
codon
gRNA
DOG1
1bp
indel
100 bp
B
Nipponbare
reference
tga5_B2
Nipponbare
reference
tga5_B2
Nipponbare
reference
tga5_B2
C
Nipponbare
reference
Nipponbare
reference
Nipponbare
reference
Fig. S12. Rice tga5 loss-of-function mutant plants contain a single base pair insertion resulting in a premature stop codon in exon 4.
